# Anthropogenic-driven loss of an adaptive radiation reduces thermal response diversity

**DOI:** 10.64898/2026.07.07.736981

**Authors:** Sophie Moreau, Bernhard Wegscheider, Dario Josi, Damien Bouffard, Martin Schmid, Timothy J. Alexander, Oliver Selz, Ole Seehausen, Conor Waldock

**Affiliations:** Independent Research Group Biodiversity, Ecosystems, and Society, Max Planck Institute for Biogeochemistry, Jena, Germany; Aquatic Ecology and Evolution, Institute of Ecology and Evolution, University of Bern, Bern, Switzerland; Department of Fish Ecology and Evolution, EAWAG, Swiss Federal Institute for Aquatic Science and Technology, Kastanienbaum, Switzerland; Wyss Academy for Nature at the University of Bern, Bern, Switzerland; Surface Waters - Research and Management, Eawag, Swiss Federal Institute of Aquatic Science and Technology, Kastanienbaum, Switzerland; Faculty of Geosciences and Environment, Institute of Earth Surface Dynamics, University of Lausanne, Lausanne, Switzerland; Aquatic Consulting, GmbH, Bern, Switzerland; Federal Office for the Environment (FOEN), Aquatic Restoration and Fisheries Section, 3011 Bern, Switzerland; Department of Zoology, School of Natural Sciences, Trinity College Dublin, Dublin, Ireland

**Keywords:** Response diversity, Climate change resilience, Adaptive radiation, Realised thermal niche, Community assembly, Eutrophication, Ecological resilience mechanism, Lake biodiversity, Ecological legacy effects

## Abstract

Biodiversity is predicted to stabilize ecosystems if species have different ecological niches that shape responses to environmental change. How this response diversity is influenced by ecological and evolutionary processes remains poorly understood. We determine the drivers of thermal response diversity of 16 Swiss peri-alpine lake-fish communities. We report the first evidence that evolutionary diversification of lineages through adaptive radiation can increase the response diversity of an ecosystem. In-situ diversification increases response diversity in the cold-deep lake environment, but non-endemic and non-native species contributed only weakly to response diversity. The loss of endemic species during historical anthropogenic eutrophication led to a negative legacy on present thermal response diversity in cold and deep lake strata. Overall, the interplay of evolutionary diversification, ecological assembly and anthropogenic impacts drives variation in response diversity. Conserving and restoring processes that generate diversity may help maintain the response diversity that underpins ecosystem stability during the Anthropocene.

## Introduction

How biodiversity contributes to ecosystem stability is critical to understand given the rate of biodiversity loss and potential for ecosystem collapse under human-induced global change (Donohue *et al*. 2016; Loreau & de Mazancourt 2013). If declines in some species can be buffered by population increase in other species, then ecosystems functions can be stabilized against environmental change (Loreau *et al*. 2021; Yachi & Loreau 1999). This community-scale “portfolio” or “insurance” effect is contributed by biodiversity if different species in a community have asynchronous responses to an environmental change. Asynchronous temporal responses of species is often approximated using by quantifying the interspecific differences in environmental responses of ecological performance proxies across spatial gradients (fitness, abundance, occupancy or demographic rates) and is often termed response diversity (Polazzo *et al*. 2024; Ross *et al*. 2023; Ross & Sasaki 2023). Response diversity has been observed to stabilize ecosystem functions in both experimental (Isbell *et al*. 2015; Leary & Petchey 2009; de Mazancourt *et al*. 2013) and observational systems (Dee *et al*. 2016; Sasaki *et al*. 2019; White *et al*. 2023). However, there still exists a very limited understanding of patterns in response diversity amongst communities in nature. Furthermore, there is a strong focus on the consequences of response diversity for ecosystem stability (Sasaki *et al*. 2019; White *et al*. 2023) but how response diversity is gained and lost due to ecological, evolutionary and anthropogenic processes has received less attention (Ross & Sasaki 2023).

Multiple ecological and evolutionary mechanisms - and human modification of these – have the potential to alter an ecosystem’s response diversity. For example, niche evolution through evolutionary adaptation of species to different environments or to each other, will generate differences in species’ environmental responses. From a set of species in the regional pool which have evolved different niches, a local community is assembled based on their environmental tolerances. This environmental filtering process determines the sets of species’ environmental responses within in a local community. If evolutionary processes cause strong ecological divergence, such as during adaptive radiations and ecological speciation, then response diversity may be particularly high (Rundle & Nosil 2005). For example, it has been hypothesised that temperature plays a role in facilitating and reinforcing speciation, so adaptive divergence towards different thermal regimes could contribute to overall response diversity (Keller & Seehausen 2012). In contrast, response diversity may be lower in communities with strong phylogenetic niche conservatism (Losos 2008). In this situation, species occupy similar abiotic niches and exhibit similar responses to environmental change (Wiens *et al*. 2010). In addition to these evolutionary processes, ecological processes that lead to more diverse communities may increase response diversity (Doak *et al*. 1998; Ives & Carpenter 2007), for example, in larger and more productive habitats that support higher species richness (Craven *et al*. 2018; Macarthur & Wilson 1967). Additionally, where local communities have access to larger regional pools response diversity can be higher (Loewen and Vinebrooke 2016). Here we fill an important knowledge gap by investigating how ecological and evolutionary processes generate response diversity in the first place.

Alongside ecological and evolutionary processes, human influences now strongly modify freshwater communities’ diversity, structure and function (Albert *et al*. 2021; Reid *et al*. 2019). An ecosystem responses to environmental change can be influenced by human modifications (Laliberté *et al*. 2010; Moore *et al*. 2010; Moore & Olden 2017). In Swiss lakes, eutrophication has led to the loss of almost a quarter of the endemic *Coregonus* species in the second half of the 20th century (Vonlanthen *et al*. 2012). Elevated nutrient inputs caused deep-water oxygen depletion which inhibited egg development of these deep-water spawning species that generally have cool-water affinity (Alexander & Seehausen 2021; Müller & Stadelmann 2004). We therefore investigate whether the loss of endemic species through eutrophication has resulted in a loss of response diversity differently in deep and shallow portions of lakes. Such context-dependent impacts would highlight how past anthropogenic influences modify the potential for an ecosystem response to additional global change drivers, such as climate warming (Jeppesen *et al*. 2012; Kraemer *et al*. 2021; Woolway *et al*. 2020). Additionally, human-assisted introduction of non-native species and translocations of species between lakes may influence response diversity. The effect of non-native species on response diversity may depend on why niches differ between native and non-native species. If non-native species share temperature requirements of native species but fill an ecological opportunity in other niche dimensions, they may have an agnostic effect on response diversity (Shea & Chesson 2002).

However, non-native species can be expected to contribute positively to response diversity, for example, if the species with establish as non-natives generally have broader environmental niches than native species (Kadye & Booth 2020). Empirical evidence shows non-native species generally exhibit high anthropogenic affinity (Jeschke & Strayer 2006), so when they replace lost native diversity, this can benefit response diversity of ecosystem functions (Moore & Olden 2017). As such, non-native species potentially have highly context dependent influences on response diversity which remain understudied. This intersection between evolutionary divergence, historical anthropogenic change, biotic exchange and thermal responses has rarely been investigated either theoretically or in real world systems.

The Swiss peri-alpine lakes provide an excellent model system to study ecological, evolutionary and anthropogenic drivers of response diversity. Here species have diverged in situ, taking advantage of empty ecological niches, resulting in independent rapid adaptive radiations within all major lakes (Alexander & Seehausen 2021). In deep oligotrophic peri-Alpine lakes, endemic species of the genera *Coregonus*, *Salvelinus*, and *Cottus* have evolved through independent adaptive radiations within multiple lakes (De-Kayne *et al*. 2022; Doenz *et al*. 2018; Hudson *et al*. 2011; Lucek *et al*. 2018; Selz *et al*. 2020; Selz & Seehausen 2023), with nearby biogeographically identical but shallower lakes exhibiting no adaptive radiation. This system therefore provides timely insights on the poorly understood contributions of ecological and evolutionary mechanisms underpinning response diversity in nature. While much work on response diversity has occurred in laboratory experiments or mesocosms, here we investigate how ecological and evolutionary processes contribute to response diversity in a real-world system of 16 lakes and explain how and why response diversity varies. We explore the relationship between thermal response diversity, species richness and past eutrophication. We assess how groups of different evolutionary and biogeographic origins contribute to variation in thermal response diversity. We do not expect species richness to directly translate into response diversity but instead we predict response diversity to be determined by community composition, environmental filtering and niche occupation. We predict endemic species to contribute unique thermal responses/niches to the community. These endemic species are of post-glacial adaptive radiations and diversified into novel deep-water niches that also have different thermal regimes. We expect this contribution to response diversity to be lost through eutrophication-driven extinctions. We test whether non-native species contribute to response diversity, perhaps rescuing response diversity lost through eutrophication, but we expect a high degree of context-dependence.

## Methods

### Analysis overview

We quantified thermal response diversity in fish communities in 16 Swiss lakes by analysing 11,449 standardized fish abundance samples and combining these with depth-specific estimates of lake temperature data. The thermal responses of 52 species were estimated using statistical models of abundance along temperature gradients. We calculated response derivatives at each temperature and used these to calculate community-level metrics of response diversity. We examined how response diversity varied across lakes and between species with varying biogeographic origins, varied across lake species richness, and lake’s eutrophication history and consequence species’ extinctions history.

### Fish community data

We analysed standardized sampling of fish communities from data collected during “*Projet Lac*”, a large project that assessed fish diversity in peri-alpine lakes (Alexander & Seehausen, 2021; Figure S1). For full description of data handing see Supporting Information 2. Fish communities along depth gradients were sampled in late summer to early autumn using standardized gillnet and electro-fishing protocols (Figure S2). The sampling protocols provided standardized sampling across the full depth range of 16 large lakes in Switzerland and were designed to standardise effort based on lake. The individuals caught in one set of nets or electrofishing section defined a single observation for a location. In total, the data consisted of 4,239 unique observations of 52 fish species (Figure S1). On average, there were 265 sampling events per lake and a total of 11,449 observations of species abundance from which to build models. Across Swiss lakes the genera Coregonus, Salvelinus and Cottus have undergone adaptive radiations with distinct species/ecomorphs between lakes having convergent ecomorphological, habitat and functional characteristics (Alexander & Seehausen 2021; Lucek *et al*. 2018; Selz *et al*. 2020; Selz & Seehausen 2023; Vonlanthen *et al*. 2012) these species with similar ecomorphology were pooled into species groups in our analysis (see Table S3).

We defined three biogeographic groups of species as “*widespread native”* (n=29) which occur across Europe and are not endemic to Switzerland, “*in-situ diversification” (n=15)* as those from the genera Coregonus, Salvelinus and Cottus which arose through post-glacial adaptive divergence to novel ecological opportunities in deep lake ecosystems starting around 12,000 years ago, and “*non-native*” (n=8) species are introduced to Switzerland. Table S3 gives an overview of nomenclature and biogeographic classification of species.

### Temperature data

Lake temperatures were simulated with the “Simstrat” model of thermal stratification version 2.1.2 (Gaudard *et al*. 2019; Råman Vinnå *et al*. 2020). Simstrat is a one-dimensional (1D) physical model for simulating the thermal structure of lakes and reservoirs, resolving the vertical dimension and assuming horizontal homogeneity. Simstrat uses lake surface area, bathymetry, meteorological and hydrological time series to simulate temperature, stratification, and ice cover in the vertical dimension. For the present study, simulations were performed with a vertical resolution of 0.5 m and a daily time resolution. Depth and time of recorded fish abundances were then matched to simulated temperatures to assign a temperature to each fish record (hereafter simply “temperature”). To aggregate over potential mismatch between daily temperature simulations and the moment of fish capture, we used the average temperature over 7 days preceding the sample observation as an indicator of the thermal habitat experienced by fish during capture.

### Modelling species’ temperature response curves

We quantified species’ realized thermal niches by modelling the response of abundance to temperature. We fitted generalized additive mixed effects models (GAMMs) for each of the 52 fish species using abundance as a response variable and temperature as a predictor (Figure S3). We fitted GAMMs with error distributions that matched species’ abundance (see Supporting information 2). We also used random intercepts to account for the difference in mean abundance between sampling (i.e., survey method) and cover the differences between lakes that are not incorporated as separate covariates due to low replication of lakes (i.e., ecological differences between lakes). We used a k of 3 which balanced curves non-linearity with physiologically realistic unimodal or sigmodal thermal responses (Clarke 2017). All GAMMs were modelled using the *mgcv* package (Wood 2011) and validated using *DHARMa* package (Hartig 2026). We compared our temperature-only models to models that also included depth as an explanatory variable of species abundance, using a k=3 for sample depth (Figure S4). While these models performed marginally better than temperature only models, we found they were often biologically implausible, so we preferred to work with the biologically plausible response curves from the temperature-only regressions to assess thermal response diversity (see Supporting Information 2 for further explanation).

### Community metrics of thermal response diversity

We calculated thermal response diversity from the variation in species responses to temperature. We measured the instantaneous responses of species characterised by the first derivative of the (non-linear) slope between abundance and temperature, at a given temperature (Figure 1; following Ross et al. 2023). To assess the variation in thermal responses at a given temperature, we simulated the full temperature gradient in Swiss lakes and made predictions of abundance for each species given the temperature (Figure 1a and 1b). We calculated the change in scaled abundance (0-1) per 0.1°C change in temperature by calculating the forward finite difference (ffd) as:

**Figure 1.**
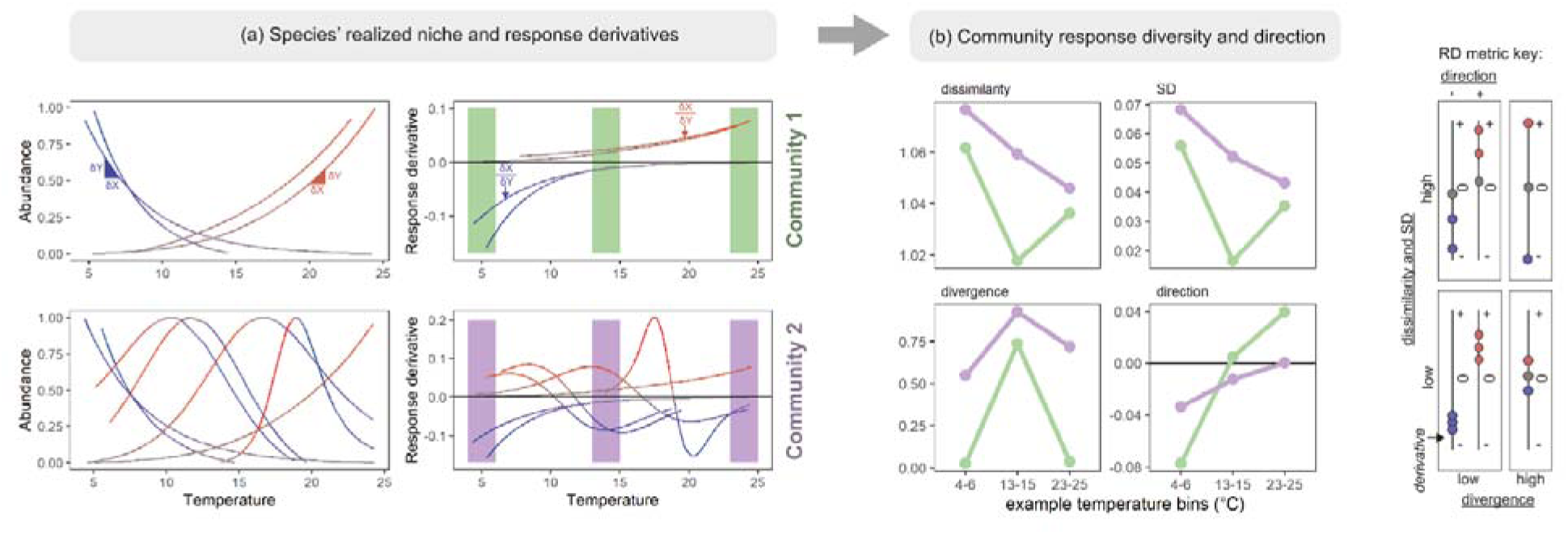
Conceptual figure mapping from species realized niches to community response diversity for two hypothetical communities with low (community 1) and high (community 2) response diversity. We estimate species’ abundance response to temperature using GAMMs (a), from which we calculate response derivatives (forward finite differences, δy / δx) across the temperature gradient (b). We the calculate the community response diversity in temperature bins (here 2°C) estimating three metrics of response diversity and one of response direction (schematically shown in the RD metric key). The overall response diversity can vary between communities, and there are also differences across temperature gradients. Our example communities are comprised from 8 selected species in our analysis as examples to demonstrate potential differences in aggregate response diversity given differences in richness, realized thermal niche shape, and community thermal structure. The colour code of species lines indicates the response derivative value. Note that we rescaled predicted abundance between 0-1.

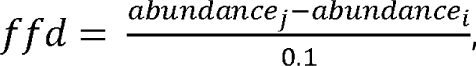, where *i* indicates abundance at current temperature and *j* indicates abundance after a 0.1°C increase in temperature (Figure 1c and 1d; Figure S8). This approximation of a derivative from predicted values was a simple, fast and intuitive method that could be applied across models with different error distributions.

From these response derivatives, we calculated three metrics of response diversity and one summary of net response direction (i.e., positive or negative net community response). From the response derivatives of each species, we calculated community scale response “dissimilarity” and “divergence” following Ross et al. (2023), as well as simply the standard deviation of response derivatives. These metrics were then calculated at a given 0.2°C temperature interval by summarising all species response derivatives at that temperature.

Dissimilarity and standard deviations quantify the overall difference in species responses. Dissimilarity indicates how differently species respond to a given environmental condition and is assessed based on a similarity-scaled metric of diversity based on the pairwise Euclidean distances in the response derivatives between all pairs of species in a community (Leinster & Cobbold 2012). The relative abundance was excluded in the calculation of dissimilarity (q=0). Dissimilarity is lowest (1), when all species respond identically and highest when all species respond differently. Note that the dissimilarity and standard deviation metrics were highly correlated (Pearson’s correlation = 0.97).

Divergence captures the ability of a community to compensate negative responses with neutral or positive ones, as classically conceptualised as the “portfolio effect” (Schindler *et al*. 2015). That is, if some species respond positively while others respond negatively, the overall community abundance can be maintained by increases counteracting decreases. Divergence was calculated following Ross et al. (2023), giving a divergence is 0 when all response derivatives are in the same direction (i.e., all positive or all negative). If response derivatives span across zero, divergence is > 0 and < 1. Divergence takes its maximum value of 1 when the absolute values of minimum and maximum response derivatives are equal.

For each lake, we constructed potential local communities at each temperature and for every 0.2°C temperature bin in each lake, we then calculated the community response diversity from the set of species’ response derivatives.

### Identifying biogeographic, environmental and anthropogenic correlates of thermal response diversity

We first assessed the patterns of metrics of response diversity and direction across thermal gradients for each lake. Next, we averaged the response diversity and direction metrics across the entire thermal gradient, giving a single response diversity and direction metric for each lake. We used spearman’s rank correlation to test how the metrics of response diversity and direction were correlated to the richness of the entire lake as well as the species richness of each biogeographic class (widespread native, in-situ diversification, non-native). Next, we assessed the same relationships but within four thermal bands within each lake (2.5 to 7.5, 7.5 to 12.5, 12.5 to 17.5 and 17.5 to 22.5) because different biogeographic groups dominate in each of these thermal bands.

Finally, we assessed how historic eutrophication of lakes related to response diversity and direction in each temperature band. We used the maximum historic phosphorus levels of each lake from Vonlanthen et al. (2012). The phosphorous data measure the annual mean total phosphorous values per lake and we take the maximum total phosphorous concentration experience by each lake across time.

### Using simulations to identify species contributions to response diversity

We simulated how species and whether they were “in situ diversification” (ISD), “non-native”, and “widespread native” contributed to community response diversity (RD). We built one simulation that contrasts ISD with widespread species and another that contrasts non-native species with widespread species. For each of these two simulations, we generated subsampled communities to standardize group sizes amongst lakes and thermal stratum. Per lake and per 0.2 degree thermal bins, we randomly subsampled the full lake-temperature strata community to 10 species. From these 10 species, we randomly dropped one single species in each of the biogeographic categories (giving two communities of 9 species). We calculated our 4 response diversity metrics for the remaining 9 species in the community. Comparing the RD of the subsampled communities indicates the effect of dropping the specific type of species. We tracked which species was dropped. We repeated this process 10 times per lake and thermal bin.

We analysed the RD from these two simulations (ISD and non-native) using mixed effects models fitted using lmerTest (version 3.2-1) package in R. We first pre-processed our data aggregating the 0.2 degree bins to four thermal bands as above (2.5 to 7.5, 7.5 to 12.5, 12.5 to 17.5 and 17.5 to 22.5). We averaged RD across lake, replicate, species biogeographic category, and species’ identity. We fitted a mixed effect model with a three-way interaction between lake, thermal strata and species biogeographic category. We used random effects of species identity and the interaction between lake and replicate within lake (to account for replicates within lakes). We used a type III Anova design and therefore tested for the marginal effects of each factor in our model. This analysis was performed on the lakes with: i) at least 1 species in each of the categories of in-situ diversification OR non-native and widespread native and ii) having species derivatives in all 4 thermal bins defined above. We therefore removed lakes Geneva, Lugano, Maggiore and Morat for ISD analysis and Lugano and Joux for the non-native analysis.

## Results

### Patterns of species’ realized thermal niches

We found that species with different biogeographic origins had qualitatively different thermal niches (Figure 2). Species originating from in-situ diversification in lakes tended to have cooler thermal niches compared to widespread species, most of which had warmer affinities. Non-native species had a range of thermal niches from cold to warm affinity. Our models of species’ thermal niches, used to derive response diversity metrics, explained a high proportion of deviance in abundance (median = 57%, IQR = 46 - 71%).

**Figure 2.**
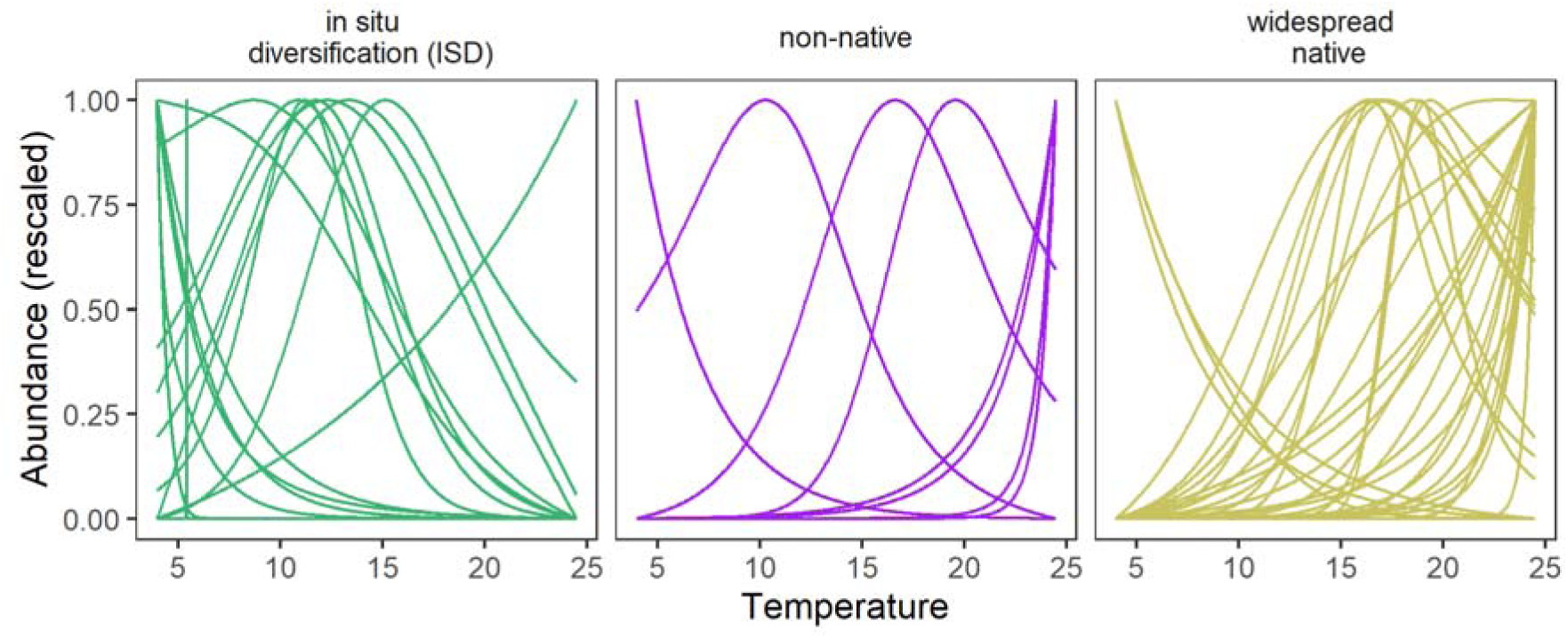
Realized thermal niches of species in different biogeographic groups. Thermal niches were identified using generalized additive models of species’ abundance modelled with temperature 7 days prior to netting. Panels indicate the biogeographic groupings relative to Switzerland peri-alpine lakes indicating “in-situ diversification” (n=15) of Salmonidae, Coregoninae, and Cottidae (sub)families, “non-native” (n=8) species originating outside of European biogeographic areas and “widespread native” species that did not originate from diversification within peri-alpine lakes (n=29). Abundance is rescaled to vary between 0 and 1 for visualisation only.

### Patterns in community response diversity within and across lakes

We found consistent patterns of response diversity across temperature gradients in lakes, with distinctive features and non-linearities along this gradient likely driven by shifting community composition along thermal gradients (Figure 3a-d). However, different metrics had different patterns of response diversity across the thermal gradient. The direction of species’ responses was generally positive, indicating warming temperatures increases net community abundance, with the strongest positive responses above 20°C. In the very coolest temperatures (<6°C) responses became increasingly negative with increased temperature.

**Figure 3.**
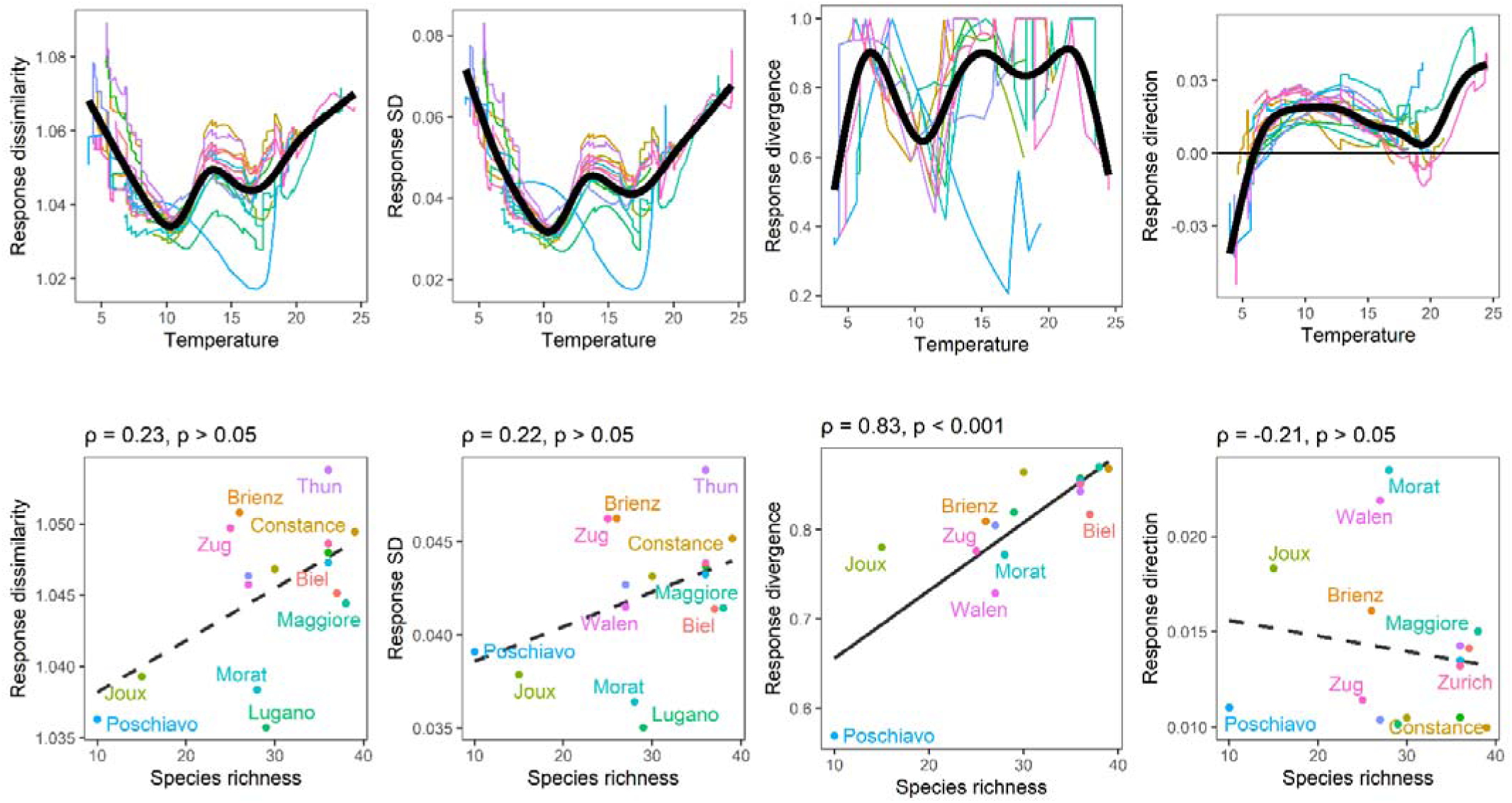
Patterns in community response diversity for within-lake temperature and across-lake richness gradients for 16 lakes. In panels a-d, lines indicate individual lakes with response diversity calculated every 0.2°C . Black lines indicate the average patterns across all lakes (produced using local polynomial regression for visualisation). In panels e-g, points indicate each lake and lines indicate linear relationships between total lake richness and response diversity with statistical summaries presented as spearman’s rank correlation estimates and significance values.

Across the entire set of lakes, total species richness had weak effects on response diversity and direction, with the exception of response divergence (Figure 3e-h). Response divergence showed a strong positive association with the overall species richness (Spearman’s rank estimate = 0.83; p < 0.001; Figure 3g).

### Richness effects are mediated by biogeographic grouping and thermal context

The weak influence of whole-lake richness on whole-lake response diversity masked a strong context dependence of the effect of richness on response diversity. We found strong relationships between richness and response diversity within specific biogeographic groups and temperature strata of lakes (Figure 4). Most strikingly, we found a strong positive relationship between species richness and response dissimilarity and response SD for species that diversified in situ at cold temperatures (Figure 4a). However, no such relationship existed for widespread native or non-native species at cold temperatures, but instead, widespread species contributed to response diversity in warmer waters (Figure 4a). The exact richness-response diversity relationships depended on the response diversity metric, with richness increasing response divergence but only for widespread native species at the warmest temperatures, and non-native and introduced species at intermediate temperatures (Figure 4b). We found no instances of significant negative richness-response diversity relationships, but a tendency that greater non-native and introduced species richness reduced response dissimilarity and SD, likely due to such species contributing non-novel thermal responses near to the community mean response (therefore lowering variance in responses).

**Figure 4.**
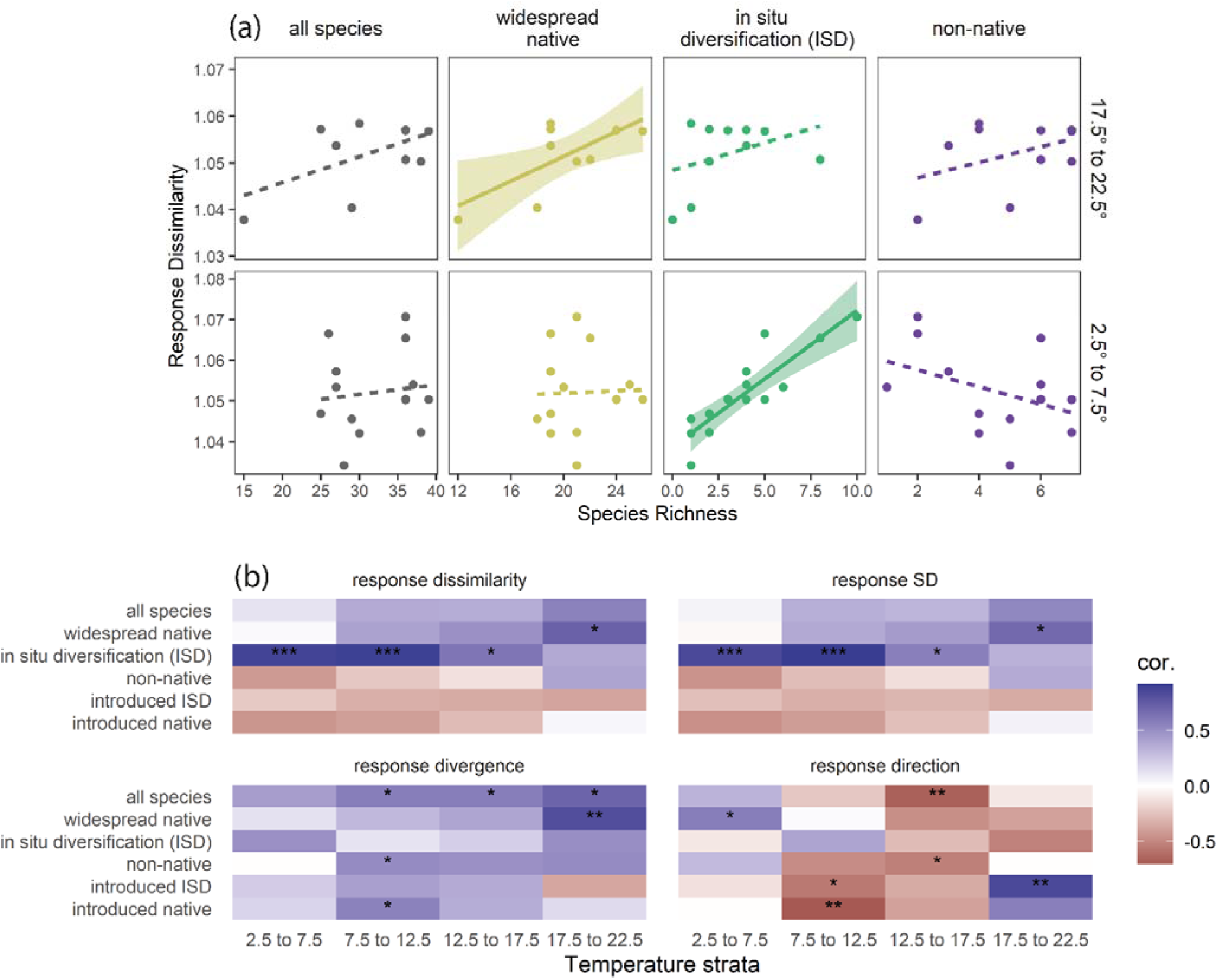
Context dependent richness-response diversity relationships depending on biogeographic origins and temperature strata. Panel a shows an example of the context dependence, whereby a positive relationships between richness and response dissimilarity only exists in specific temperature strata for species groups. Significance of relationships is determined by Spearman’s rank correlations (solid = p<0.05; dashed = p>0.05). Lines represent linear relationships fitted with least squares with shaded areas indicating 95% confidence intervals. Panel b shows the matrix of biogeographic groups and temperature strata indicating negative (red) to positive (blue) richness-response diversity relationships. The significance of richness-response diversity Spearman’s rank estimates (ρ) are indicated as p>0.05 = empty, p < 0.05 = *, p < 0.01 = **, p < 0.001 = ***.

### Drivers of response diversity identified through simulations

We found that removing in-situ diversification species significantly reduced response diversity when compared with removing widespread species from a standardised community (Table S4). The biogeographic origin of species alone explained 10% of the variation in response divergence but only 2% of response dissimilarity. The effect of removing in-situ diversification species differed across thermal strata (Table S4) and was especially strong at 12.5 to 17.5°C for all metrics (Figure 5a-c). The effect of removing in-situ diversification species differed across different lakes for response divergence (Figure 5e) but not for response dissimilarity (Figure 5f). While the three-way interaction between lake, temperature strata and biogeographic origin was highly significant (Table S4), the largest effects were through two-way interactions between biogeography and temperature strata and lake level differences (Table S4). Almost all in-situ diversification species have net positive contributions to response dissimilarity (Figure S9) but many native species also contribute positively, with a fraction of common native species having negative or neutral net contributions (e.g., *Barbus barbus*, *Lota lota* and *Squalius cephalus* and *Salvelinus umbla*).

**Figure 5.**
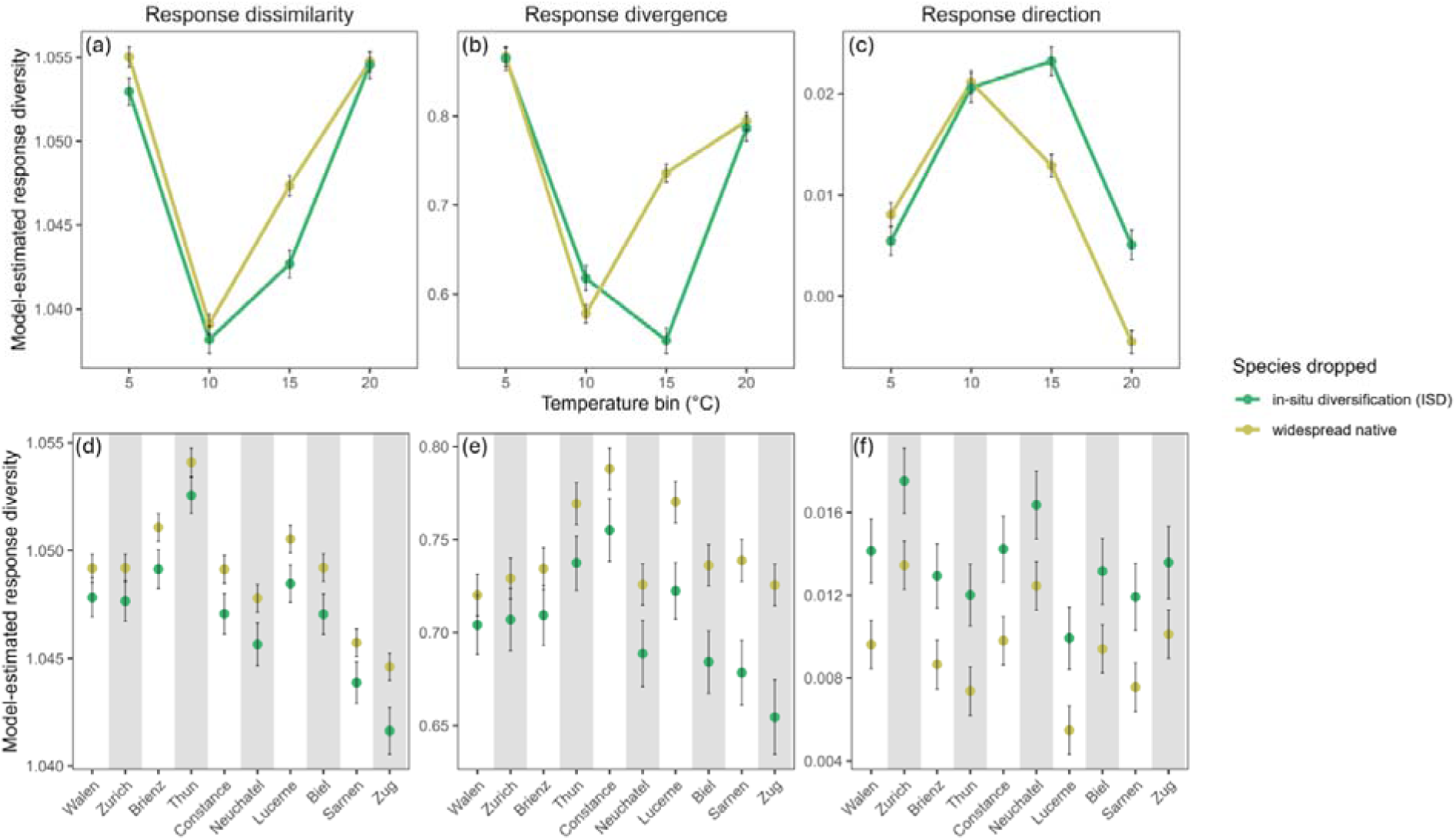
Species’ contribution to response diversity across thermal gradients, biogeographic groups and lakes as indicated by a dropping one random species in each focal group from a standardized community. Estimated marginal mean values ± 95% CI from an ANOVA design shows the effect of each grouping, thermal strata and lake on response diversity metrics as indicated on the y-axis. The panels a-c show estimate values across temperature strata. The panels d-f show estimates across each lake. Colours indicate when in-situ diversification (green) are dropped from the community compared with widespread native species (yellow). For the equivalent plot for non-native species see Figure S10.

We tested the above patterns for non-native species compared with widespread species. The influence of removing non-native species was weaker than removing in-situ diversification species (Table S4), and while the interaction between biogeography and thermal strata were significant, the effect sizes were 0.07-0.18x the effect of in-situ diversification species (Figure S10, Table S4). Underlying these net effects were both positive and negative contributions to response diversity of non-native species (Figure S11).

### Eutrophication reduced response diversity in cold lake strata

We found a negative relationship between response diversity and the strength of historic eutrophication, with statistical significance only in the coldest thermal strata between 2.5°C to 7.5°C (Figure 6). At this temperature strata, we found significant negative relationships for response dissimilarity (ρ = -0.57, p<0.05), divergence (ρ = -0.74, p < 0.01) and negative non-significant relationships for response SD (ρ = -0.51, p>0.05) and no correspondence with response direction (ρ = 0.02, p>0.05). Further, there was a tendency for response diversity to negatively relate to the strength of historic eutrophication at 7.5°C to 12.5°C but none of these associations were significant (Figure 6). The strength of these relationships then weakened between 12.5°C to 22.5°C indicating no evidence eutrophication history influenced response diversity in the warmer thermal strata despite effects in cold strata (Figure 6).

**Figure 6.**
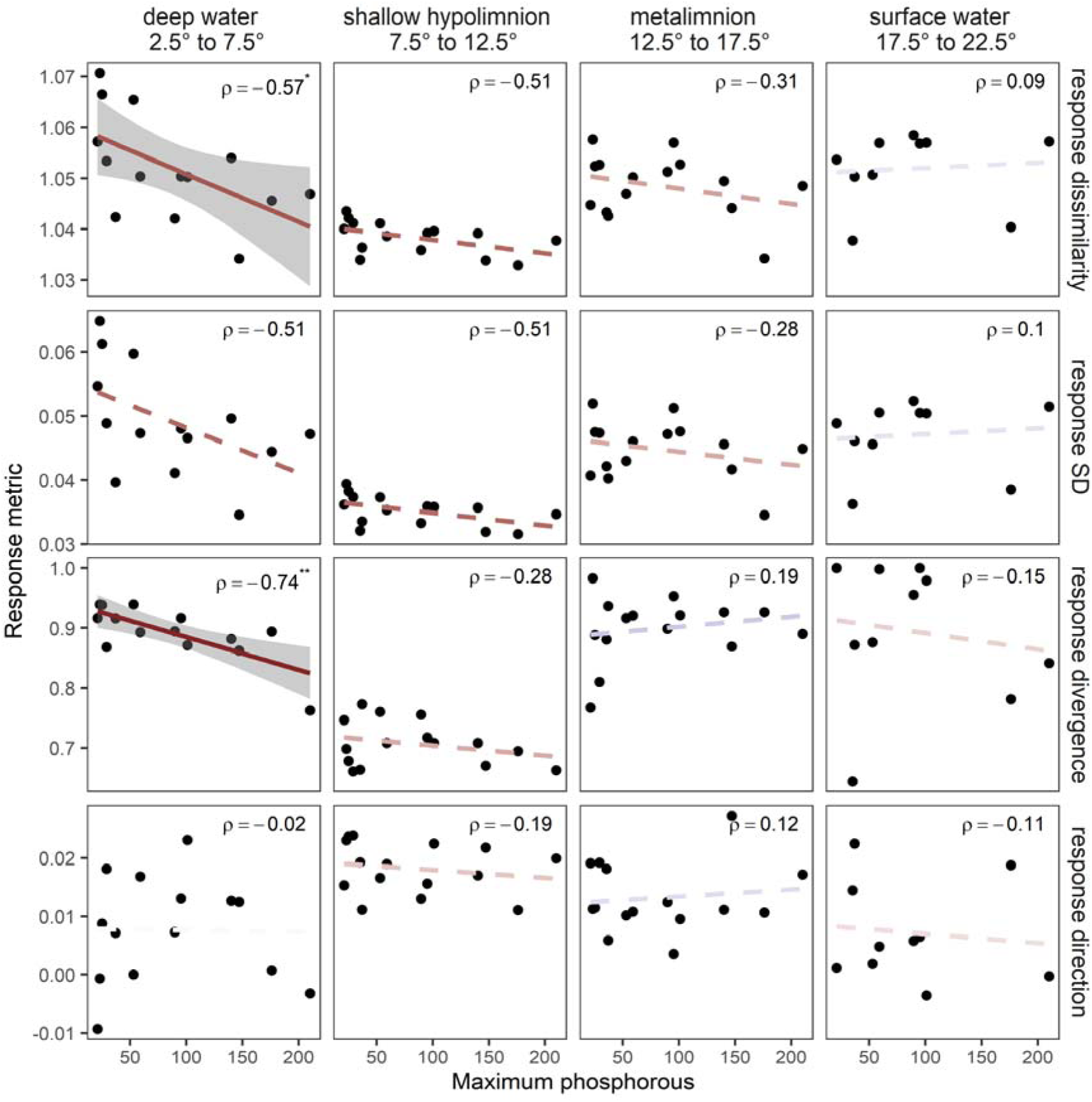
Lake historic eutrophication reduces response diversity in cold lake strata. Points indicate values of maximum historic phosphorus levels and response diversity metrics for all species present in given temperature strata. Lines represent linear relationships fitted with least squares with shaded areas indicating 95% confidence intervals. Significance of relationships is determined by Spearman’s rank correlations (solid = p<0.05; dashed = p>0.05). Line colour visualises negative (red) to positive (blue) relationship. The significance of phosphorus-response diversity Spearman’s rank estimates (ρ) are indicated as p>0.05 = empty, p < 0.05 = *, p < 0.01 = **, p < 0.001 = ***.

## Discussion

Factors that influence the gain and loss of response diversity demand research attention given the role of response diversity contributing to the stability of ecosystem functions (Ross & Sasaki 2023). We discovered that evolutionary processes, thermal gradients in the environment, community assembly and historic anthropogenic eutrophication interact to determine the variation in response diversity within and between lakes. The positive relationship we found between species richness and response diversity varied across thermal strata because of the different compositions of species assembling and diversifying in these different thermal strata in the last 12,000 years. In more recent decades, anthropogenic eutrophication caused deoxygenation below the thermocline which differentially impacted species dependent on well-oxygenated, cold and profundal thermal strata for egg development (Vonlanthen *et al*. 2012). These species contributed to response diversity through their adaptation towards complementary thermal niches (Keller & Seehausen 2012). Accordingly, the loss of deepwater endemic species also meant the loss of their unique contributions to response diversity, potentially modifying the stability of the system. In contrast, eutrophication in the warmer shallow waters did not lead to species loss. The non-endemic species assembled here through immigration and made weaker contributions to response diversity than the lineages diversifying in adaptive radiations. Thus, within a single lake, fundamentally different ecological and evolutionary processes shaped response diversity in the deep-cool and shallow-warm regions. Quantifying this context-dependence contributes a deeper understanding of drivers of variation in response diversity. Richness was not the sole driver of response diversity, but instead a lake’s community composition is an important contributor to response diversity. This means we must move beyond richness-stability relationships and instead consider how niche diversification contributes to response diversity and the potential consequences for stability (Hooper *et al*. 2005; Ives & Carpenter 2007; Xu *et al*. 2021).

Our work indicates that understanding the evolutionary origins of species richness is necessary to understand variation in response diversity. We report the first instance of diversification through adaptive radiation having contributed to response diversity, to our knowledge. The evolution of ecological diversity through adaptive radiation in peri-alpine lakes, especially in *Coregonus*, has filled vacant ecological niches post-glaciation. In this system, the partitioning of reproductive, phenological, and trophic niche space in sympatry (Doenz *et al*. 2018; Vonlanthen *et al*. 2012) has led to an associated partitioning of (realized) thermal niche space (as observed in Kelly *et al*. 2015). The steep thermal gradients in this system may reinforce ecological speciation leading to fundamental thermal niche diversification but this process is very poorly explored (Keller & Seehausen 2012). Given that speciation reversal has occurred in this system – whereby sympatric species are now less ecologically and genetically distinct due to increased gene-flow, or have completely collapsed into a single population (Vonlanthen *et al*. 2012) – we likely currently observe a weaker effect of adaptive radiation on response diversity than would have been present historically. This inference is supported given that consistently oligotrophic lakes, representing an unimpacted state, had higher response diversity than lakes that were recently eutrophic.

Species originating from adaptive radiations contributed more strongly to response diversity compared with widespread native and non-native species. This contrast likely reflects different evolutionary processes whereby niche divergence in sympatry facilitates diversification. In contrast, widespread or non-native species diversified in allopatry outside the deep peri-alpine lakes under genetic drift and with weak, and not generally divergent, thermal selection. Our results support that under allopatry we expect more redundant thermal niches compared to sympatry which drives complementary thermal niches (Keller & Seehausen 2012), especially if divergence is scaled over evolutionary time. As such, phylogenetic divergence does not relate to response differences in our system even though widespread species cover a far wider diversity of branches along the teleost evolutionary tree. In other contexts, phylogenetic divergence does covary with response differences and communities have higher ecological stability (Cadotte *et al*. 2012), and phylogenetically distinct non-native species can enhance response diversity when establishment traits align with response traits (Moore & Olden 2017). Interestingly, there exists phylogenetic structure in thermal traits of freshwater fishes but this did not manifest as contributing strongly to response diversity in our work (Comte & Olden 2017). For example, we might have expected coexisting species with deep evolutionary divergences to contribute to response diversity, but these lineages likely diverged along non-thermal niche axes (e.g., diet, habitat) which allows co-existence in more similar thermal habitats during post-glacial community assembly (Alexander & Seehausen 2021). As such, the covariance between response traits and phylogenetic distances, and how diversification processes contribution to this covariance, appear an important consideration in understanding variation in response diversity.

More broadly, our findings hint at an underappreciated role of the geography of speciation and the ecology of reproductive isolation in contributing to response diversity. Different taxonomic groups often diversify through fundamentally different processes which then shapes sets of species’ response differently. As such, there exists evolutionary constrained variation amongst taxonomic groups in their potential response diversity, as seen here. We should perhaps expect different levels of response diversity (for a given species richness) when contrasting lineages emerging from allopatric speciation cycles (e.g., birds and riverine fishes; Dias *et al*. 2013; Sholihah *et al*. 2021; Tobias *et al*. 2020), and with those emerging through interspecific interactions causing prezygotic reproductive isolation (e.g., plants; Baack *et al*. 2015), with those diversifying into vacant niches (sympatric adaptive radiation; Miller 2021; Seehausen & Wagner 2014). This variation in evolutionary processes, structured by evolutionary lineage, biome and geography, may generate fundamentally different hypotheses of how response diversity contributes to stability of ecosystem functions and ecosystem service change under global change. As yet, little work exists comparing how the process of evolutionary diversification, and specifically adaptive radiation, contributes to ecological stability mechanisms like response diversity, and such hypotheses still remain poorly conceptualised with little empirical or theoretical work (Ross & Sasaki 2023). Our case study thereby shines an important light on the relevance of evolutionary processes in understanding global change impacts on ecosystems through response diversity (Brodersen & Seehausen 2014; Faith *et al*. 2010).

Our work highlights that context dependent biodiversity change is expected if one global change stressor modifies the potential responses to a second. Here, we find that eutrophication has led to a reduction in the potential thermal responses specifically for deep-cold lake fish communities. Such eutrophication driven loss of thermal response diversity may have consequences for how climate change impacts these communities. For example, even though the bottom waters are expected to warm slower than the surface waters (Råman Vinnå *et al*. 2021), the profundal ecosystem has lost response diversity, which may modify future climate change responses in eutrophic lakes (and also lakes that have transiently been eutrophic but are oligotrophic now). This link between transient anthropogenic eutrophication and climate change sensitivity is entirely contingent on the adaptive processes leading to niche diversification in these specific lakes and the specific way that eutrophication differentially impacts deeper waters through deoxygenation. A similar phenomenon occurs in the Laurentian Great Lakes, whereby an invasive species *Alosa pseudoharengus* is expected to have reduced the abundance and diversity of the native Coregonus species complex, with an expected reduction in the biological portfolio and resilience of lake food webs (Hale et al. 2026). Nature in the Anthropocene is defined by multiple co-occurring stressors acting together (Bowler *et al*. 2020), but the influence of one stressor constraining and influencing the set of potential responses to other stressors is rarely considered. More often, multi-stressor impacts are conceptualised as interactions between stressors that amplify or dampen their impacts (Birk *et al*. 2020). Future work should also focus on how ecological and evolutionary processes interact with stressors (here eutrophication) to modify the potential responses to another stressor (here climate change).

Our quantification of species ecological niches using observational data limits us to quantify the realized, rather than fundamental, thermal niche of species. Realized niche estimates are potentially confounded by unobserved variables, for example depth, oxygen and light, which can shift niche estimates when included alongside temperature in our niche models (Figure S4). However, most thermal niche estimates were very similar, or actually far less realistic (e.g., *Cottus gobio* Rhine, *Cobitis bilineata*, *Squalius cephalus*), when also including depth as a covariate due to the behaviour of multiple regressions under highly correlated variables (Morrissey & Ruxton 2018). As such, we used the single variable models and interpret results in the knowledge that multiple environmental gradients underpin the spatial distribution of fish in lakes. As a further caveat, we sampled during a short temporal window in just one year per lake, which was (coincidentally) near the period of highest within-year thermal variation along depth gradients.

We note our metric of realized niche likely integrates both population demographic processes and behavioural thermoregulation, whereby individuals move to preferred thermal habitats along depth gradients (Magnuson et al. 1979; Megner et al. 2010; Raby 2020). Translating response diversity to community stability from realized niche metrics that represent behavioural regulation may poorly map to the processes driving community dynamics. In general, although the biological realism of realised niches have been challenged (Lee-Yaw *et al*. 2021), empirical evidence suggests strong correspondence can exist between aquatic organisms’ fundamental niches and realised niches (Lv *et al*. 2026; Payne *et al*. 2021; Rezende & Bozinovic 2019). As such, we likely recover an important index of thermal tolerance from our realized niche estimates that, while imperfect, helps us understand the drivers of thermal response diversity within a community.

We also note that the insurance hypothesis framework, by which response diversity should relate to ecosystem stability, was developed for time-series responses of species’ populations to environmental change (Yachi & Loreau 1999; Loreau et al 2021). We therefore make a space-for-time assumption, common in global change ecology, whereby population abundance across spatial thermal gradients is assumed to predict temperature related changes through time (Lovell et al. 2023). Our dataset contains most of the largest peri-alpine lakes in Switzerland, and despite the somewhat limited sample size of only 16 lakes, we recover theoretically expected and mechanistically logical results. In addition, it remains a question for future work to address whether the variation in response diversity observed here has a real impact on the stability of these lake fish communities, which would require time-series data to test explicitly.

In conclusion, this work represents an important first step in understanding how the biodiversification process, rather than just biodiversity per-se, contributes to response diversity with consequences for the expected change in ecosystems under environmental change. We find strong context dependencies that will likely also exist in other systems in determining variation in the gains and losses of response diversity. By unpacking how ecological and evolutionary processes contribute to variation in response diversity, our work contributes important insights to explaining the high amounts of inherent variability in biodiversity change and ecosystem stability in the Anthropocene (Blowes *et al*. 2019; Hillebrand *et al*. 2020; Johnson *et al*. 2024).

## Acknowledgements

This work is part of the project LANAT-3 ‘Den Biodiversitätsverlust der Gewässer stoppen — trotz Klimawandel’ funded by the Wyss Academy for Nature through the implementation programme with the canton of Bern (Office for Agriculture and Nature) and by the Federal Office for the Environment (FOEN). This project funding supported C.W., B.W., and D.J. Thanks to the Aquatic Ecology and Evolution group at the University of Bern and the EAWAG Department of Fish Ecology & Evolution for thoughtful discussions and feedback. We acknowledge the use of generative AI (Claude, Opus 5.5) to support code development.

## Statement of Authorship

*S. M., C. W., D. J. and T. A., and curated the data with expertise in lake temperature modelling, lake biodiversity sampling, and species taxonomy provided by M. S., D. B., O. Selz and O. Seehausen. Feedback during project development was provided by all authors. The idea and methodology was developed by S.M, C.W. and B.W. Supervision was provided by C.W, B.W and O. Seehausen. The investigation was carried out by S. M., C. W., B. W, D. J and O. Seehausen. The formal analyses, software development, data validation and visualisation were led by S.M. and C.W. The resources were provided by O. Seehausen. The original draft was written by S.M. and C.W. and all authors contributed to the review and editing of the manuscript. All authors approved the final version for submission*.

## Data Accessibility Statement

The full code repository to reproduce the scripts are provided at https://github.com/cwaldock1/lake-response-diversity/releases/tag/v1, these scripts and data are also provided at figshare with the private link https://figshare.com/s/7a7f865dca7aa28a8ad1. This will be made available as a full DOI at https://doi.org/10.6084/m9.figshare.32407596 on acceptance.

