## Supporting Information 1 for "Anthropogenic-driven loss of an adaptive radiation reduces thermal response diversity"

**Supplementary information**

Table S1. Summary of number of species and number of community samples per lake and sampling protocol.

| **Lake** | **Protocol** | **n. species** | **n. observations** |
| --- | --- | --- | --- |
| Biel | CEN | 20 | 78 |
| Biel | VERT | 15 | 25 |
| Biel | electro | 19 | 39 |
| Brienz | CEN | 12 | 60 |
| Brienz | VERT | 12 | 48 |
| Brienz | electro | 7 | 15 |
| Constance | CEN | 23 | 207 |
| Constance | VERT | 24 | 201 |
| Constance | electro | 21 | 77 |
| Geneva | CEN | 18 | 189 |
| Geneva | VERT | 17 | 299 |
| Geneva | electro | 16 | 79 |
| Joux | CEN | 6 | 27 |
| Joux | VERT | 8 | 25 |
| Joux | electro | 6 | 22 |
| Lucerne | CEN | 16 | 95 |
| Lucerne | VERT | 18 | 112 |
| Lucerne | electro | 12 | 29 |
| Lugano | CEN | 12 | 51 |
| Lugano | VERT | 15 | 60 |
| Lugano | electro | 8 | 20 |
| Maggiore | CEN | 22 | 132 |
| Maggiore | VERT | 20 | 75 |
| Maggiore | electro | 11 | 58 |
| Morat | CEN | 16 | 45 |
| Morat | VERT | 17 | 42 |
| Morat | electro | 9 | 24 |
| Neuchatel | CEN | 20 | 137 |
| Neuchatel | VERT | 21 | 85 |
| Neuchatel | electro | 11 | 20 |
| Poschiavo | CEN | 6 | 48 |
| Poschiavo | VERT | 5 | 59 |
| Poschiavo | electro | 3 | 24 |
| Sarnen | CEN | 19 | 85 |
| Sarnen | VERT | 14 | 20 |
| Sarnen | electro | 11 | 37 |
| Thun | CEN | 22 | 89 |
| Thun | VERT | 20 | 73 |
| Thun | electro | 12 | 22 |
| Walen | CEN | 15 | 77 |
| Walen | VERT | 14 | 69 |
| Walen | electro | 9 | 26 |
| Zug | CEN | 16 | 52 |
| Zug | VERT | 17 | 96 |
| Zug | electro | 13 | 29 |
| Zurich | CEN | 19 | 71 |
| Zurich | VERT | 20 | 120 |
| Zurich | electro | 15 | 45 |

Table S2. Summary of number of abundance estimates per species as well as the minimum, maximum, median and mean abundance of those samples.

| **Species** | **n. observations** | **min. abun.** | **max. abun.** | **median abun.** | **mean abun.** |
| --- | --- | --- | --- | --- | --- |
| Salvelinus namaycush | 3 | 1 | 1 | 1 | 1 |
| Salvelinus sp. Limnetic | 5 | 1 | 1 | 1 | 1 |
| Alosa fallax | 8 | 1 | 4 | 2 | 2 |
| Carassius gibelio | 8 | 1 | 3 | 2 | 1.6 |
| Ameiurus melas | 9 | 1 | 6 | 2 | 2.4 |
| Salvelinus sp. | 9 | 1 | 4 | 1 | 1.8 |
| Anguilla anguilla | 11 | 1 | 5 | 1 | 2.1 |
| Cottus gobio Rhine | 12 | 1 | 6 | 2 | 2.9 |
| Alburnus arborella | 14 | 1 | 25 | 2 | 4.6 |
| Alosa agone | 14 | 1 | 25 | 2 | 4.5 |
| Salmo sp. Blackspot | 15 | 1 | 3 | 1 | 1.5 |
| Cobitis bilineata | 18 | 1 | 2 | 1 | 1.2 |
| Micropterus salmoides | 20 | 1 | 6 | 1 | 1.8 |
| Cottus gobio unknownlineage | 21 | 1 | 11 | 1 | 2 |
| Silurus glanis | 22 | 1 | 2 | 1 | 1.1 |
| Barbatula affinisfluvicola | 26 | 1 | 34 | 2 | 4.3 |
| Squalius squalus | 29 | 1 | 7 | 1 | 1.5 |
| Coregonus sp. benthic profundal | 30 | 1 | 8 | 1 | 1.7 |
| Salvelinus sp. Profundal | 30 | 1 | 3 | 1 | 1.1 |
| Barbus barbus | 31 | 1 | 15 | 1 | 1.9 |
| Cyprinus carpio | 31 | 1 | 8 | 1 | 1.9 |
| Cottus sp. Po | 36 | 1 | 7 | 2 | 2.2 |
| Cottus sp. Profundal | 38 | 1 | 7 | 1 | 1.8 |
| Coregonus sp. large pelagic | 42 | 1 | 5 | 1 | 1.2 |
| Phoxinus csikii | 46 | 1 | 22 | 2 | 4.7 |
| Scardinius hesperidicus | 46 | 1 | 38 | 1 | 3.6 |
| Barbatula ommata | 51 | 1 | 22 | 1 | 3.4 |
| Blicca bjoerkna | 55 | 1 | 49 | 2 | 4.1 |
| Salaria fluviatilis | 68 | 1 | 49 | 2 | 4.2 |
| Coregonus sarnensis | 74 | 1 | 2 | 1 | 1.2 |
| Coregonus sp. balchen | 74 | 1 | 3 | 1 | 1.1 |
| Cottus gobio littoral | 77 | 1 | 12 | 1 | 1.6 |
| Lepomis gibbosus | 88 | 1 | 16 | 1 | 2 |
| Tinca tinca | 128 | 1 | 6 | 1 | 1.3 |
| Coregonus sp felchen | 129 | 1 | 8 | 1 | 1.5 |
| Abramis brama | 147 | 1 | 51 | 1 | 2.9 |
| Sander lucioperca | 147 | 1 | 27 | 2 | 3.1 |
| Esox spp | 161 | 1 | 4 | 1 | 1.2 |
| Salvelinus umbla | 161 | 1 | 10 | 1 | 1.5 |
| Scardinius erythrophthalmus | 167 | 1 | 17 | 2 | 3.1 |
| Salmo trutta | 196 | 1 | 26 | 2 | 3.1 |
| Gobio gobio | 201 | 1 | 58 | 2 | 4.2 |
| Squalius cephalus | 251 | 1 | 32 | 1 | 2.7 |
| Gasterosteus aculeatus | 252 | 1 | 282 | 2 | 10 |
| Lota lota | 291 | 1 | 31 | 1 | 1.7 |
| Coregonus sp. albeli | 370 | 1 | 38 | 1 | 2.1 |
| Leuciscus leuciscus | 380 | 1 | 107 | 2 | 4 |
| Gymnocephalus cernua | 467 | 1 | 200 | 3 | 9.7 |
| Alburnus alburnus | 698 | 1 | 171 | 1 | 3.5 |
| Coregonus sp. | 1325 | 1 | 40 | 1 | 2 |
| Rutilus rutilus | 1718 | 1 | 311 | 2 | 7.6 |
| Perca fluviatilis | 3199 | 1 | 590 | 4 | 17 |

Table S3. Overview of biogeographic status per species/ecotype per lake as determined by Alexander and Seehausen (2021). W = widespread native, NN = non-native, ISD = in-situ diversification.

| Species | Status | Biel | Constance | Geneva | Lucerne | Morat | Neuchatel | Sarnen | Zug | Zurich | Thun | Brienz | Walen | Joux | Lugano | Maggiore | Poschiavo |
| --- | --- | --- | --- | --- | --- | --- | --- | --- | --- | --- | --- | --- | --- | --- | --- | --- | --- |
| Abramis brama | W | x | x | x | x | x | x | x | x | x | x | x |  |  |  |  |  |
| Alburnoides bipunctatus | W | x | x |  | x | x |  |  | x | x |  | x |  |  |  |  |  |
| Alburnus alburnus | W | x | x | x | x | x | x | x | x | x | x | x | x | x |  |  |  |
| Alburnus arborella | W |  |  |  |  |  |  |  |  |  |  |  |  |  | x | x |  |
| Alosa agone | W |  |  |  |  |  |  |  |  |  |  |  |  |  | x | x |  |
| Alosa fallax | W |  |  |  |  |  |  |  |  |  |  |  |  |  |  | x |  |
| Ameiurus melas | NN |  | x | x |  |  |  |  |  | x |  |  |  |  |  | x |  |
| Anguilla anguilla | W | x | x | x | x | x | x | x | x | x | x |  | x |  | x | x |  |
| Barbatula affinisfluvicola | W |  | x | x |  |  |  |  |  |  |  |  |  |  |  |  |  |
| Barbatula ommata | W | x |  |  | x | x | x | x | x | x | x | x | x |  |  |  |  |
| Barbus barbus | W | x | x | x |  | x | x | x | x | x | x | x | x |  |  |  |  |
| Barbus plebejus | W |  |  |  |  |  |  |  |  |  |  |  |  |  | x | x |  |
| Blicca bjoerkna | W | x | x | x | x | x | x | x | x | x | x |  | x |  |  |  |  |
| Carassius gibelio | NN | x | x |  |  | x | x |  |  | x |  |  |  |  | x | x |  |
| Chondrostoma nasus | W | x | x |  | x | x | x | x |  | x | x |  | x |  |  |  |  |
| Chondrostoma soetta | W |  |  |  |  |  |  |  |  |  |  |  |  |  | x | x |  |
| Cobitis bilineata | NN | x | x | x |  | x | x |  |  |  |  |  |  |  |  |  |  |
|  | W |  |  |  |  |  |  |  |  |  |  |  |  |  |  | x |  |
| Coregonus sp large pelagic | NN |  |  |  |  |  |  |  |  |  | x |  |  |  |  |  |  |
| Coregonus sp albeli | ISD | x |  |  | x |  | x |  |  | x | x | x | x |  |  |  |  |
|  | NN | x |  |  |  |  |  |  |  |  |  |  |  |  |  |  |  |
| Coregonus sp balchen | ISD | x | x |  |  | x | x |  | x | x | x | x | x |  |  |  |  |
| Coregonus sp felchen | ISD |  | x |  | x |  |  | x |  | x | x | x | x |  |  |  |  |
| Coregonus sp | ISD |  |  |  | x |  |  |  |  |  | x |  |  |  |  |  |  |
| Coregonus sp balchen | NN |  |  | x |  |  |  |  |  |  |  |  |  |  | x | x |  |
| Coregonus sp benthic profundal | ISD |  |  |  | x |  |  |  |  |  | x |  |  |  |  |  |  |
| Coregonus sarnensis | NN |  |  |  |  |  |  |  |  |  |  |  |  |  |  | x |  |
|  | ISD |  |  |  |  |  |  | x |  |  |  |  |  |  |  |  |  |
| Coregonus sp | NN |  |  |  |  |  |  |  |  |  |  |  |  | x |  | x |  |
| Coregonus sp large pelagic | ISD |  | x |  |  |  |  |  |  |  |  |  |  |  |  |  |  |
| Cottus gobio littoral | ISD | x |  |  | x |  | x | x | x | x | x | x | x |  |  |  |  |
| Cottus sp Profundal | ISD |  |  |  | x |  |  |  |  |  | x |  | x |  |  | x |  |
| Cottus gobio Rhine | ISD |  | x | x |  |  |  |  |  |  |  |  |  |  |  |  |  |
| Cottus gobio unknownlineage | ISD | x |  |  |  |  |  | x |  |  |  |  |  |  |  |  |  |
| Cottus sp Po | ISD |  |  |  |  |  |  |  |  |  |  |  |  |  | x | x | x |
| Cyprinus carpio | W | x | x | x | x | x | x | x | x | x | x | x | x | x | x | x |  |
| Esox spp | W | x | x | x | x | x | x | x | x | x | x | x | x | x | x | x |  |
| Gasterosteus aculeatus | NN | x | x |  | x |  | x |  |  | x |  |  |  |  |  |  |  |
| Gasterosteus gymnurus | NN |  |  | x |  |  | x |  |  |  |  |  |  |  |  |  |  |
|  | W |  |  |  |  |  |  |  |  |  |  |  |  |  |  | x |  |
| Gobio gobio | W | x | x | x | x | x | x | x | x | x | x | x | x |  | x | x |  |
| Gobio obtusirostris | W |  | x |  |  |  |  |  |  |  |  |  |  |  |  |  |  |
| Gymnocephalus cernua | NN |  | x | x | x | x |  | x | x | x |  |  |  |  |  | x |  |
| Lampetra planeri | W | x | x |  | x |  | x |  |  | x | x |  |  |  |  |  |  |
| Lepomis gibbosus | NN | x | x | x | x |  | x |  | x | x |  |  |  |  | x | x |  |
| Leuciscus leuciscus | W | x | x | x | x | x | x | x | x | x | x | x | x | x |  |  |  |
| Lota lota | W | x | x |  | x |  | x | x | x | x | x | x | x | x | x | x |  |
|  | NN |  |  | x |  |  |  |  |  |  |  |  |  |  |  |  |  |
| Micropterus salmoides | NN | x |  | x |  | x | x |  |  |  |  |  |  |  | x | x |  |
| Oncorhynchus mykiss | NN | x | x |  | x | x | x | x | x | x | x | x | x | x | x | x |  |
| Padogobius bonelli | W |  |  |  |  |  |  |  |  |  |  |  |  |  | x | x |  |
| Perca fluviatilis | W | x | x | x | x | x | x | x | x | x | x | x | x | x | x | x |  |
| Phoxinus csikii | W | x | x |  | x |  | x |  |  | x | x | x | x | x |  |  |  |
|  | NN |  |  |  |  |  |  |  |  |  |  |  |  |  |  |  | x |
| Phoxinus sp | NN |  |  |  |  |  |  |  |  |  | x |  |  |  |  |  | x |
|  | W |  |  | x |  |  |  |  |  |  |  |  |  |  |  |  | x |
| Rhodeus amarus | W | x | x |  |  | x | x |  |  |  |  |  |  |  |  |  |  |
|  | NN |  |  |  |  |  |  |  |  |  |  |  |  |  |  | x |  |
| Rutilus aula | W |  |  |  |  |  |  |  |  |  |  |  |  |  | x | x |  |
| Rutilus rutilus | W | x | x | x | x | x | x | x | x | x | x | x | x | x |  |  |  |
|  | NN |  |  |  |  |  |  |  |  |  |  |  |  |  | x | x |  |
| Salaria fluviatilis | NN |  |  | x |  |  |  |  |  |  |  |  |  |  |  |  |  |
|  | W |  |  |  |  |  |  |  |  |  |  |  |  |  | x | x |  |
| Salmo cenerinus | W |  |  |  |  |  |  |  |  |  |  |  |  |  |  |  | x |
| Salmo labrax | NN |  |  |  |  |  |  |  |  |  |  |  |  |  |  |  | x |
| Salmo marmoratus | W |  |  |  |  |  |  |  |  |  |  |  |  |  | x | x | x |
| Salmo sp Blackspot | W |  |  |  |  |  |  |  |  |  |  |  |  |  |  |  | x |
| Salmo trutta | W | x | x | x | x | x | x | x | x | x | x | x | x | x |  |  |  |
|  | NN |  |  |  |  |  |  |  |  |  |  |  |  |  | x | x | x |
| Salvelinus namaycush | NN |  |  |  | x |  |  |  |  | x | x | x |  |  |  |  | x |
| Salvelinus sp Profundal | ISD |  | x |  | x |  |  |  |  |  | x |  | x |  |  |  |  |
| Salvelinus sp Limnetic | ISD |  |  |  | x |  |  |  |  |  | x | x |  |  |  |  |  |
| Salvelinus umbla | W | x | x | x | x | x | x |  | x | x |  | x | x |  |  |  |  |
|  | NN |  |  |  |  |  |  | x |  |  |  |  |  |  | x | x | x |
| Salvelinus sp | ISD |  |  |  |  |  |  |  |  |  | x |  |  |  |  |  |  |
| Sander lucioperca | NN | x | x |  | x | x | x | x | x |  |  |  |  | x | x | x |  |
| Scardinius erythrophthalmus | W | x | x | x | x | x | x | x | x | x | x | x | x |  |  |  |  |
| Scardinius hesperidicus | NN | x |  | x |  |  | x |  |  | x | x |  |  |  |  |  |  |
|  | W |  |  |  |  |  |  |  |  |  |  |  |  |  | x | x |  |
| Silurus glanis | W | x | x |  |  | x | x |  |  | x | x |  |  |  |  |  |  |
|  | NN |  |  |  |  |  |  |  |  |  |  |  |  |  | x | x |  |
| Squalius cephalus | W | x | x | x | x | x | x | x | x | x | x | x | x | x |  |  |  |
| Squalius squalus | W |  |  |  |  |  |  |  |  |  |  |  |  |  | x | x |  |
| Telestes muticellus | W |  |  |  |  |  |  |  |  |  |  |  |  |  | x | x |  |
| Thymallus thymallus | W | x | x | x | x |  | x | x |  | x | x | x | x | x |  |  |  |
| Tinca tinca | W | x | x | x | x | x | x | x | x | x |  | x | x | x | x | x |  |
|  |  | **Biel** | **Constance** | **Geneva** | **Lucerne** | **Morat** | **Neuchatel** | **Sarnen** | **Zug** | **Zurich** | **Thun** | **Brienz** | **Walen** | **Joux** | **Lugano** | **Maggiore** | **Poschiavo** |

Table S4. Assessing the effect of biogeography (endemic vs. native), thermal strata and lake on response diversity through drop-one out simulations. Analysis of variance table for the fixed effects, with random effects: (1 | species_dropped) and (1 | lake:rep_id). Response diversity metrics across lakes, temperature bins and biogeographic groups from our simulation. We included three-way interactions between all variables.

| **Response variable** | **Term** | **df** | **F** | **p** |  |
| --- | --- | --- | --- | --- | --- |
| Response dissimilarity | lake | 9, 6583.6 | 262.13 | < .001 | *** |
|  | thermal strata | 3, 7508.9 | 5,765.56 | < .001 | *** |
|  | biogeography | 1, 38.0 | 16.41 | < .001 | *** |
|  | lake × thermal strata | 27, 7493.5 | 181.88 | < .001 | *** |
|  | lake × biogeography | 9, 6583.6 | 1.59 | 0.112 |  |
|  | thermal strata × biogeography | 3, 7508.9 | 96.08 | < .001 | *** |
|  | lake × thermal strata × biogeography | 27, 7493.5 | 3.76 | < .001 | *** |
| Response divergence | lake | 9, 6210.6 | 49.55 | < .001 | *** |
|  | thermal strata | 3, 7510.0 | 3,953.05 | < .001 | *** |
|  | biogeography | 1, 38.4 | 23.50 | < .001 | *** |
|  | lake × thermal strata | 27, 7494.7 | 55.99 | < .001 | *** |
|  | lake × biogeography | 9, 6210.6 | 6.47 | < .001 | *** |
|  | thermal strata × biogeography | 3, 7510.0 | 663.35 | < .001 | *** |
|  | lake × thermal strata × biogeography | 27, 7494.7 | 11.24 | < .001 | *** |
| Response direction | lake | 9, 7217.9 | 109.88 | < .001 | *** |
|  | thermal strata | 3, 7500.5 | 5,315.50 | < .001 | *** |
|  | biogeography | 1, 34.8 | 21.23 | < .001 | *** |
|  | lake × thermal strata | 27, 7488.6 | 130.99 | < .001 | *** |
|  | lake × biogeography | 9, 7217.9 | 0.60 | 0.795 |  |
|  | thermal strata × biogeography | 3, 7500.5 | 593.07 | < .001 | *** |
|  | lake × thermal strata × biogeography | 27, 7488.6 | 3.06 | < .001 | *** |
| Response SD | lake | 9, 6518.5 | 266.95 | < .001 | *** |
|  | thermal strata | 3, 7509.1 | 6,443.54 | < .001 | *** |
|  | biogeography | 1, 37.6 | 13.47 | < .001 | *** |
|  | lake × thermal strata | 27, 7493.4 | 185.01 | < .001 | *** |
|  | lake × biogeography | 9, 6518.5 | 1.70 | 0.083 |  |
|  | thermal strata × biogeography | 3, 7509.1 | 113.24 | < .001 | *** |
|  | lake × thermal strata × biogeography | 27, 7493.4 | 4.03 | < .001 | *** |

Table S5. Assessing the effect of biogeography (non-native vs. native), thermal strata and lake on response diversity through drop-one out simulations. Analysis of variance table for the fixed effects, with random effects: (1 | species_dropped) and (1 | lake:rep_id). Response diversity metrics across lakes, temperature bins and biogeographic groups from our simulation. We included three-way interactions between all variables.

| Response variable | Effect | df | F | p |  |
| --- | --- | --- | --- | --- | --- |
| Response dissimilarity | lake | 9, 7358.3 | 137.53 | < .001 | *** |
|  | thermal strata | 3, 7367.3 | 8,260.31 | < .001 | *** |
|  | biogeography | 1, 34.0 | 1.90 | 0.177 |  |
|  | lake × thermal strata | 27, 7350.8 | 60.71 | < .001 | *** |
|  | lake × biogeography | 9, 7358.3 | 0.86 | 0.561 |  |
|  | thermal strata × biogeography | 3, 7367.3 | 17.66 | < .001 | *** |
|  | lake × thermal strata × biogeography | 27, 7350.8 | 3.42 | < .001 | *** |
| Respose divergence | lake | 9, 7364.8 | 116.43 | < .001 | *** |
|  | thermal strata | 3, 7377.1 | 3,857.53 | < .001 | *** |
|  | biogeography | 1, 34.3 | 1.47 | 0.233 |  |
|  | lake × thermal strata | 27, 7354.2 | 39.86 | < .001 | *** |
|  | lake × biogeography | 9, 7364.8 | 8.19 | < .001 | *** |
|  | thermal strata × biogeography | 3, 7377.1 | 52.93 | < .001 | *** |
|  | lake × thermal strata × biogeography | 27, 7354.2 | 6.46 | < .001 | *** |
| Response direction | lake | 9, 7364.3 | 62.82 | < .001 | *** |
|  | thermal strata | 3, 7376.4 | 4,501.62 | < .001 | *** |
|  | biogeography | 1, 34.4 | 0.68 | 0.415 |  |
|  | lake × thermal strata | 27, 7354.0 | 157.85 | < .001 | *** |
|  | lake × biogeography | 9, 7364.3 | 0.97 | 0.463 |  |
|  | thermal strata × biogeography | 3, 7376.4 | 42.14 | < .001 | *** |
|  | lake × thermal strata × biogeography | 27, 7354.0 | 3.23 | < .001 | *** |
| Respose SD | lake | 9, 7358.2 | 137.97 | < .001 | *** |
|  | thermal strata | 3, 7367.1 | 8,120.40 | < .001 | *** |
|  | biogeography | 1, 34.0 | 2.03 | 0.163 |  |
|  | lake × thermal strata | 27, 7350.7 | 63.41 | < .001 | *** |
|  | lake × biogeography | 9, 7358.2 | 1.04 | 0.404 |  |
|  | thermal strata × biogeography | 3, 7367.1 | 17.12 | < .001 | *** |
|  | lake × thermal strata × biogeography | 27, 7350.7 | 4.07 | < .001 | *** |

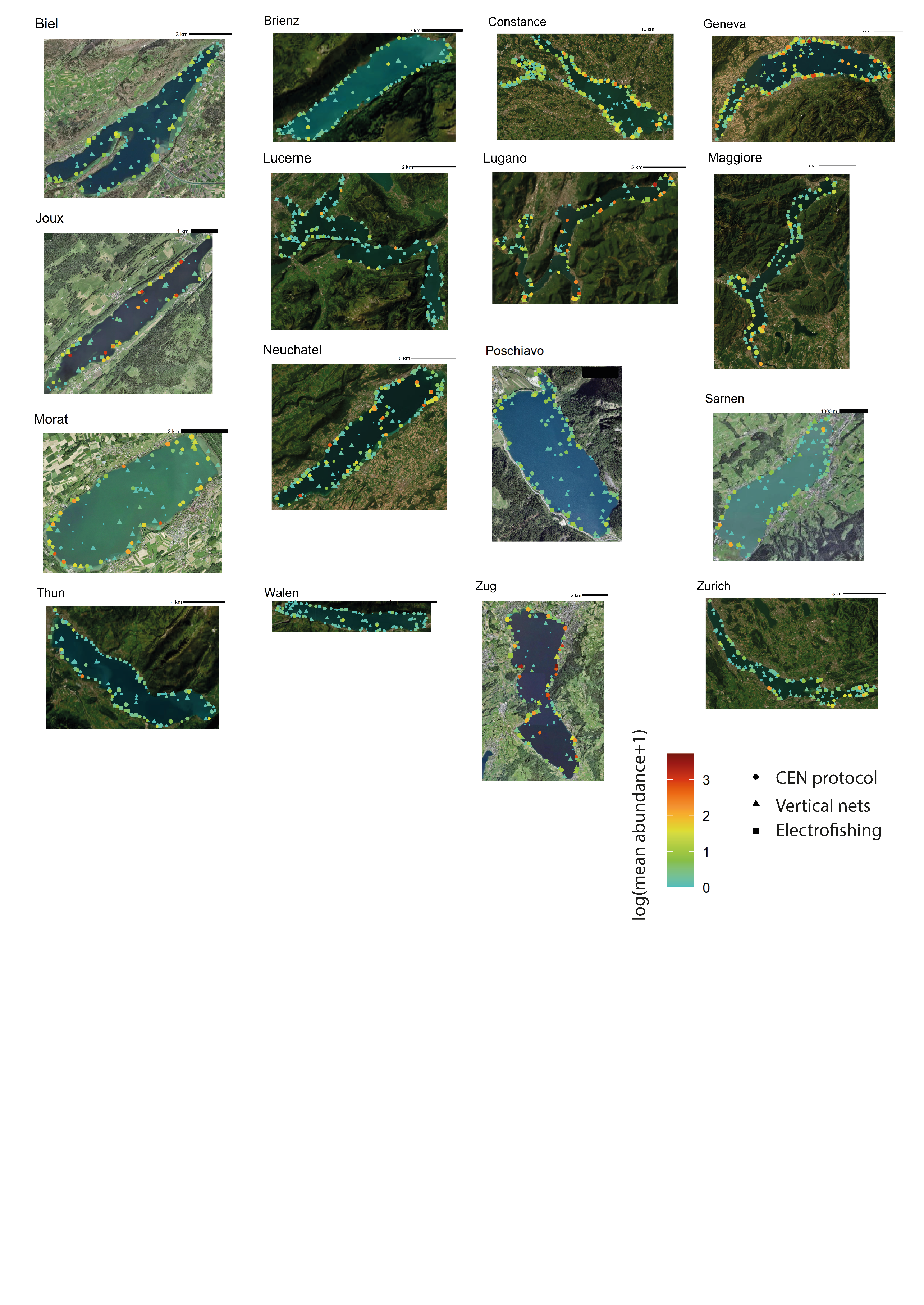

Figure S1. Spatial distribution of *projet lac* records used in analysis of abundance-temperature relationships and their corresponding sampling protocols. Size of points indicates species richness.

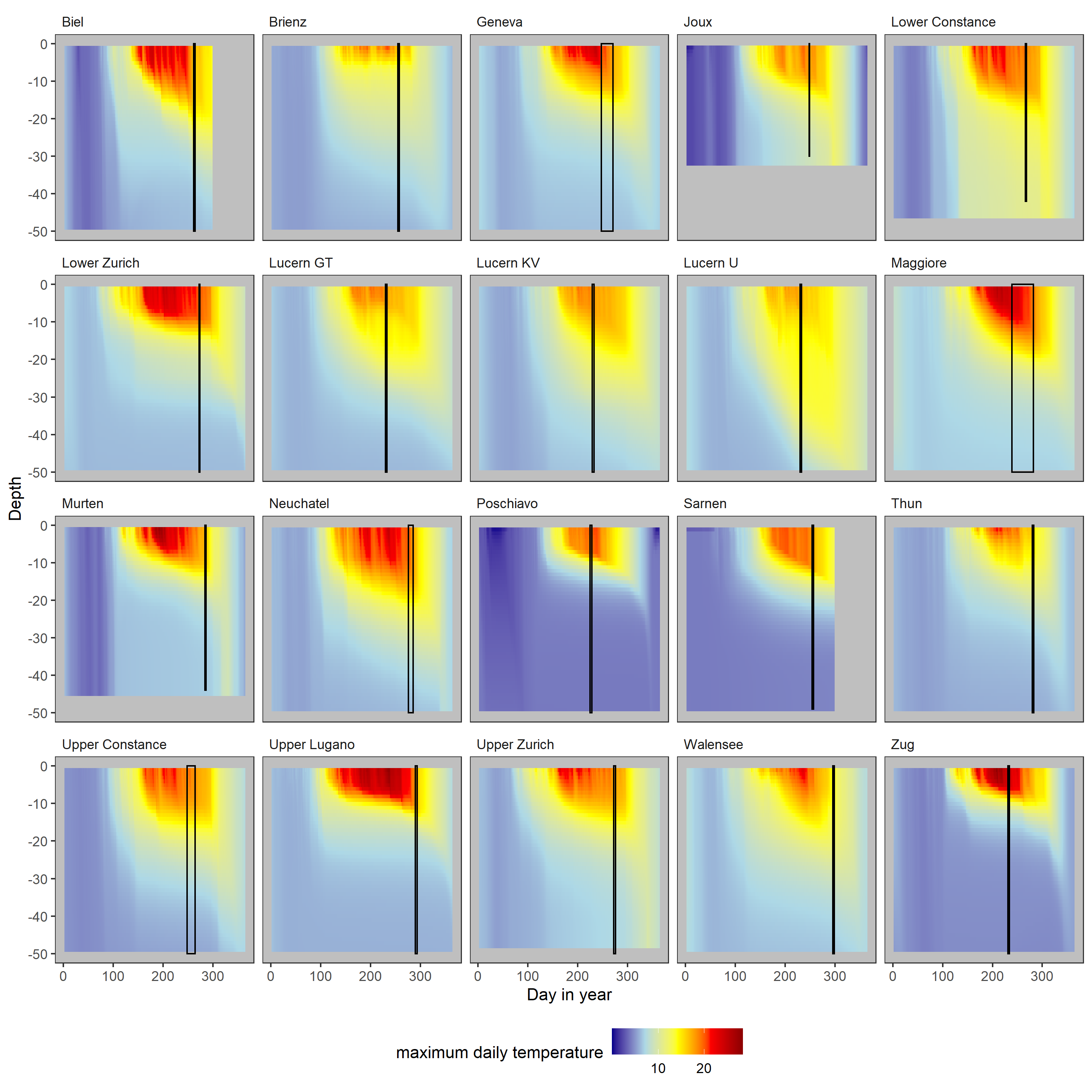

Figure S2. Temperature across the year (x-axis) and depth (y-axis) with sampling windows indicated with black boxes for the 16 lakes used in our analysis. Note that lake basins are separated here whereas they are aggregated in the main text. Colour indicates maximum daily temperature whereas in the main text we use the mean temperature 7 days prior to sampling as our index of species’ thermal environment. Note that this figure is truncated to 50m to best visualise the thermocline gradient, whereas the maximum depth of most of the lakes apart from Lakes Murten and Joux is greater than 50m.

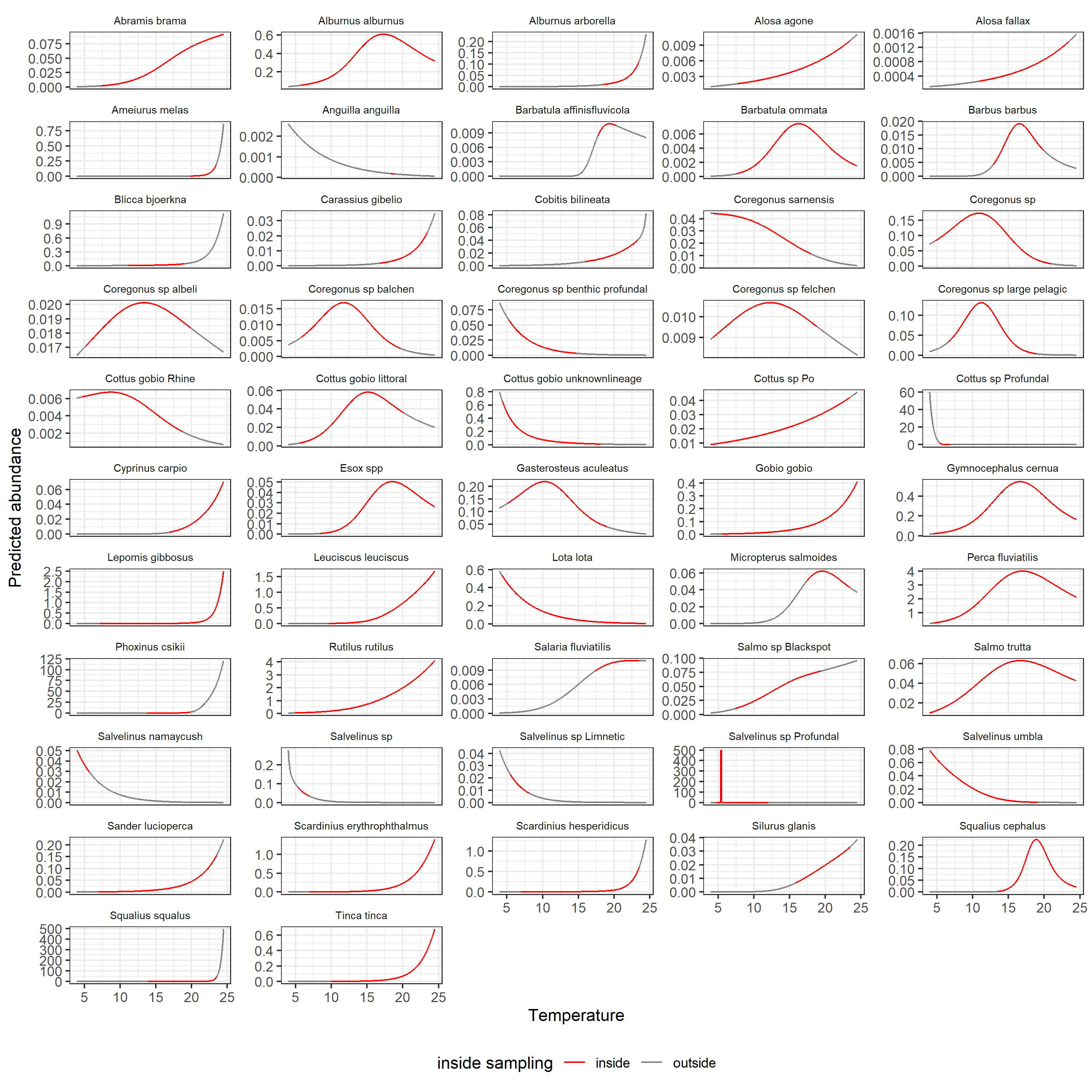

Figure S3. Abundance-temperature relationships for 52 species. Colour indicates whether predictions are inside or outside of the sampled temperature range for a given species. Note that “Salvelinus sp. Profundal” abundances are truncated to 500 as models predicted extreme abundances for this species.

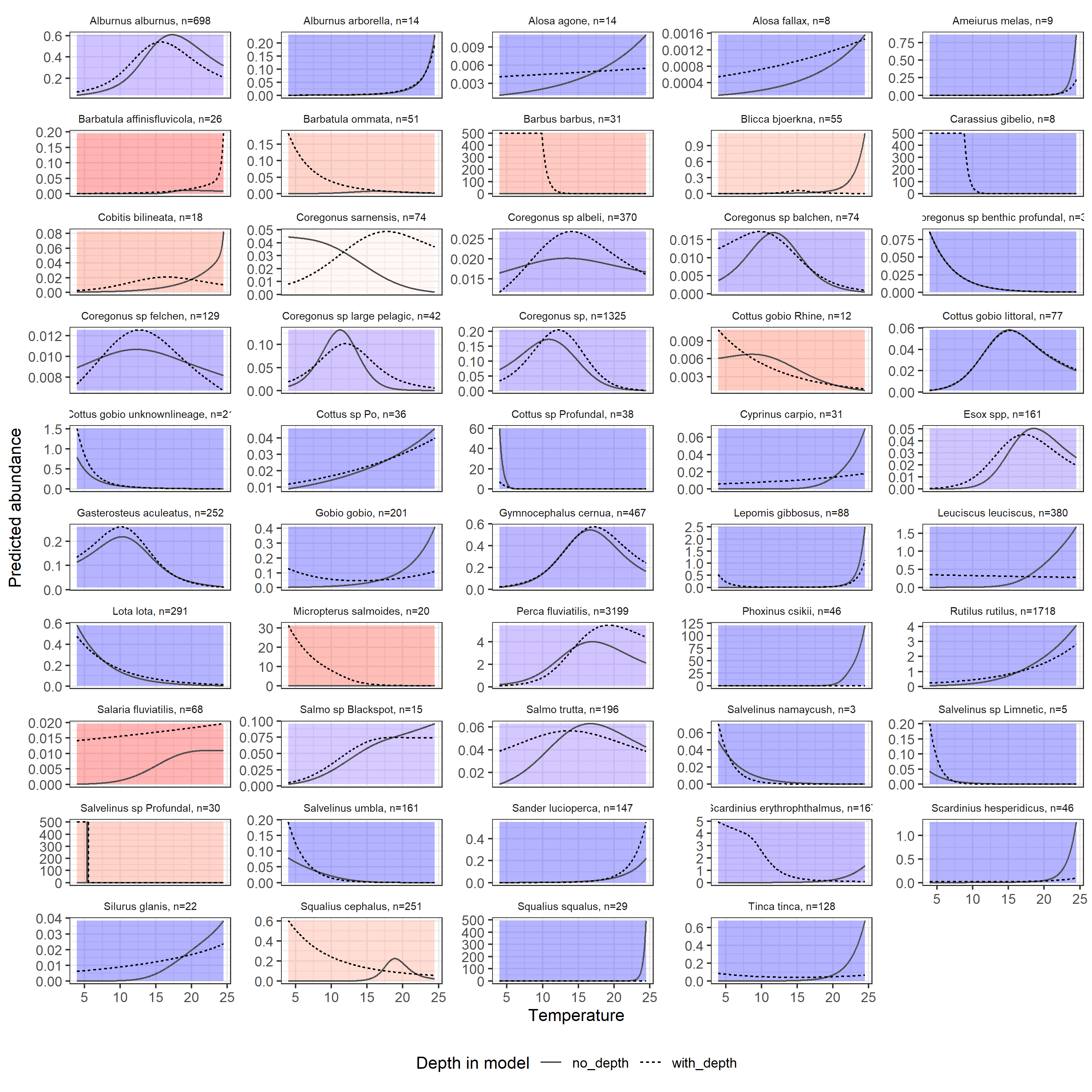

Figure S4. Comparison of response curves from GAMMs with depth included (dashed) or excluded (solid) as a covariate (with k=3) during model fitting. The amount of data per species is indicated in the facet titles. The background colour indicates the Spearman’s rank correlation between the response derivative (finite forward difference) including and excluding depth as a covariate. In 78% of cases the response derivatives showed positive correlations between models including and excluding depth, and 65% of species has rank correlations > 0.9. The mean spearman’s rank correlation across species was 0.58 and the median was 0.99. The cases where thermal response curves do not match between models generally had thermal response curves that did not match expectations based on known thermal affinities of species (e.g., *Scardinius erythrophthalmus*, *Squalius cephalus*, and *Barbatula ommata)* – likely indicating an issue with model identifiability when incorporating both depth and temperature.

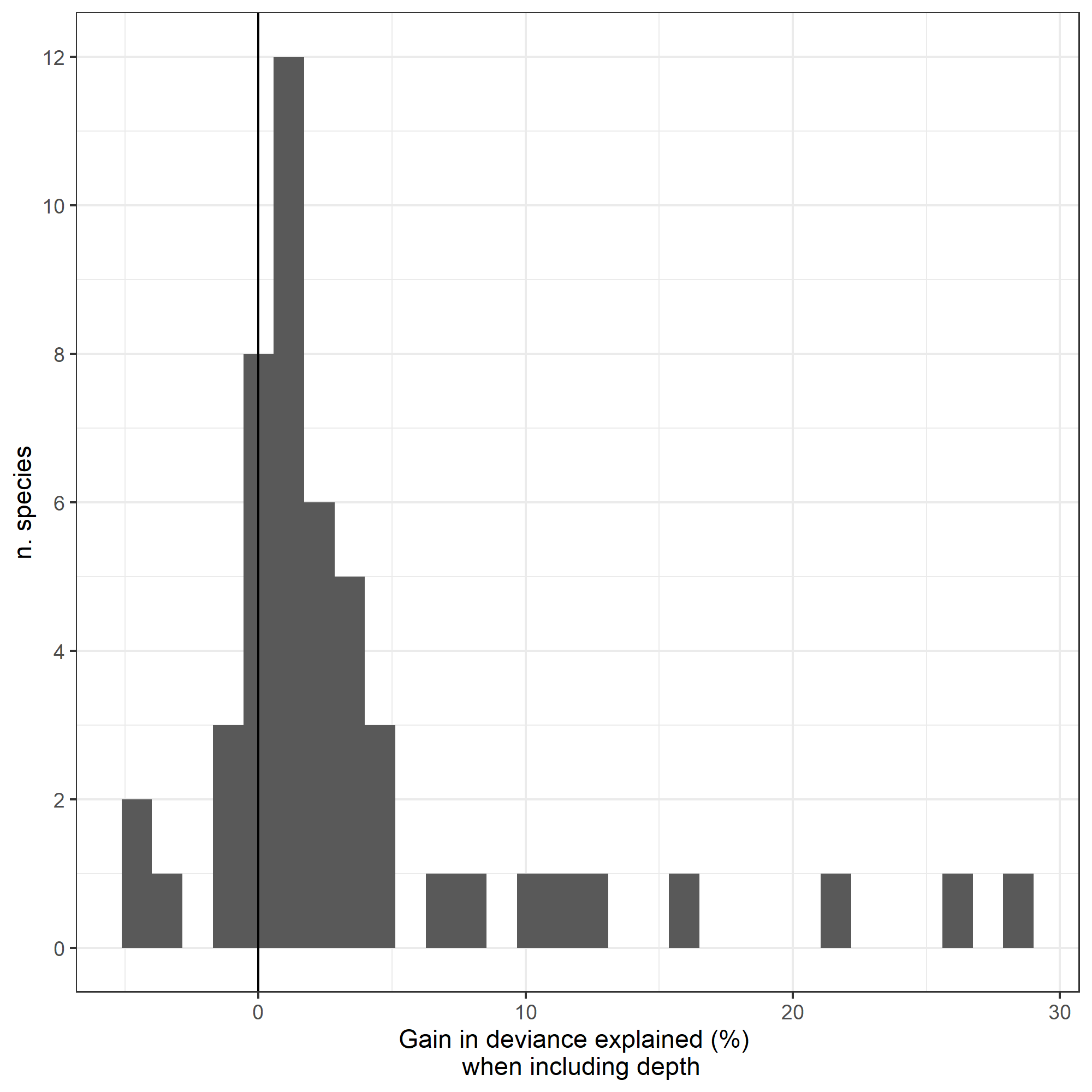

Figure S5. The difference in model deviance explained when including depth as a covariate (with k=3) during model fitting. The median gain in model performance was 1% and the mean gain 3%, the species with gains > 10% were *Lota lota* (10%), *Blicca bjoerkna* (11%), *Alosa agone* (12%), *Barbus barbus* (16%), *Micropterus salmoides* (22%), *Cyprinus carpio* (26%) and *Carassius gibelio* (28%).

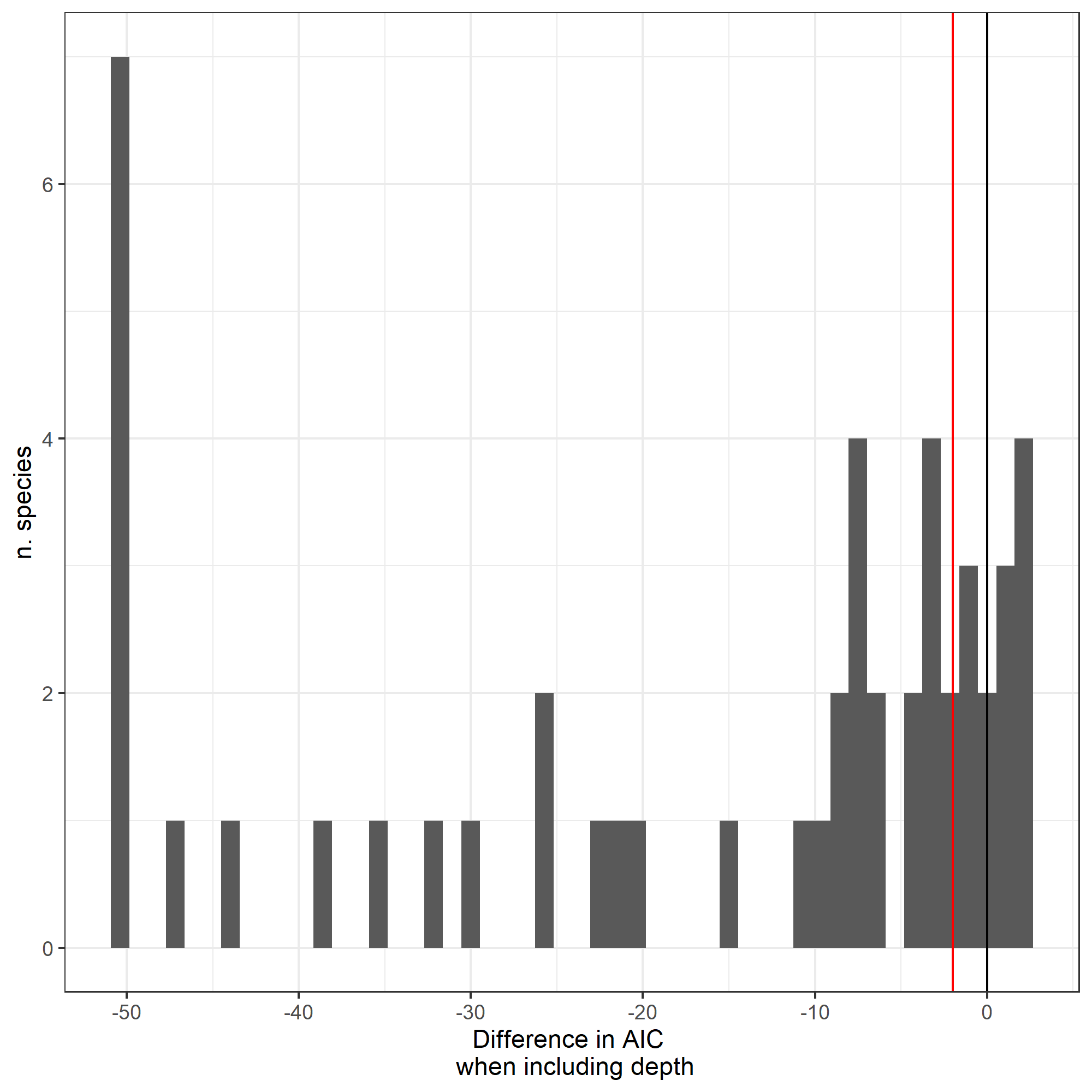

Figure S6. The difference in AIC when including depth as a covariate (with k=3) during model fitting. 36/52 species show improvements in model fit when including depth as a covariate.

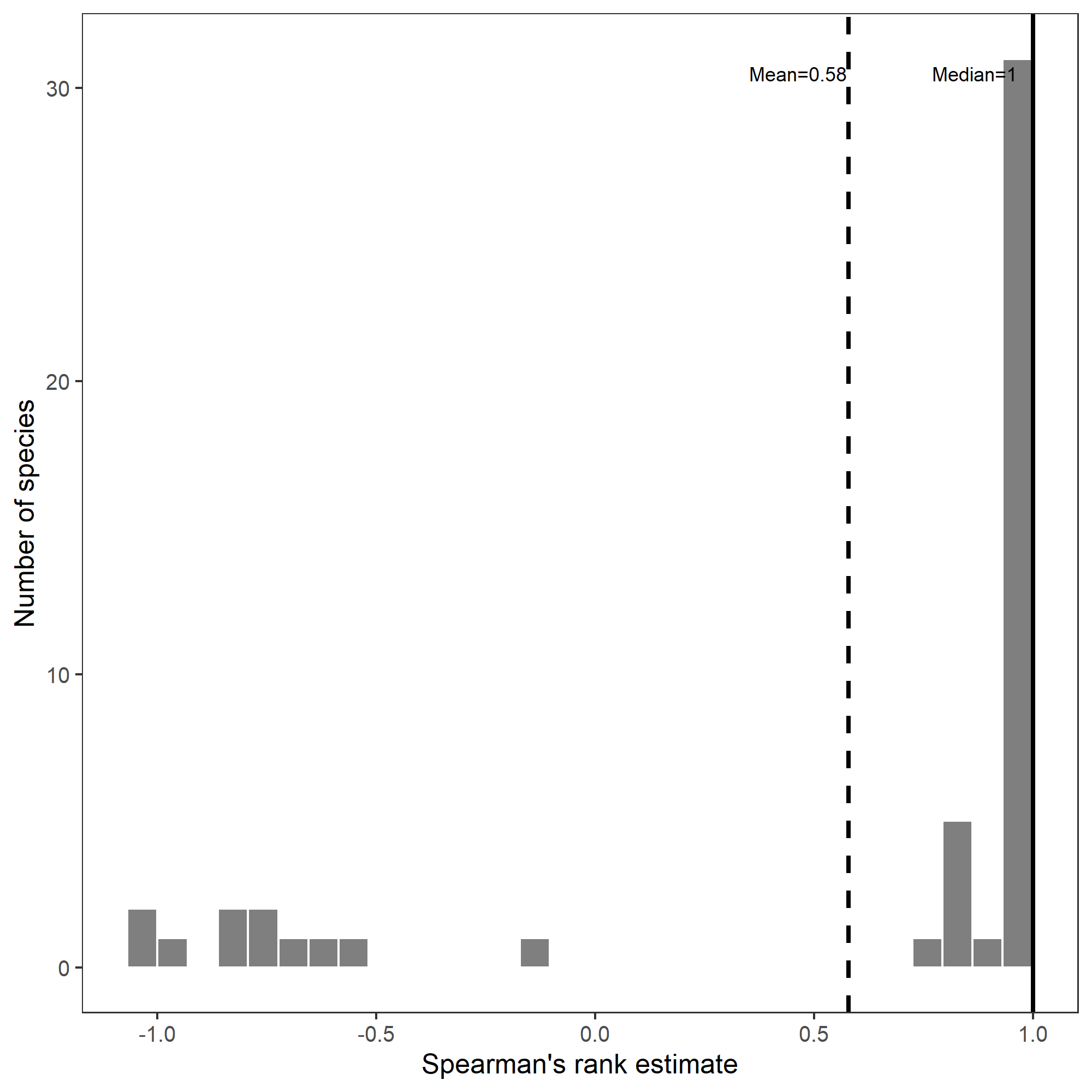

Figure S7. The Spearman’s rank correlation between response derivatives (forward finite difference) when including depth as a covariate and when excluding depth as a covariate in our thermal niche models. 78% of species had positive rank correlations, and 65% of species has rank correlations > 0.9. The species with negative correlations tended to have thermal responses, when including depth, that did not align with expectations based on the known spatial distribution and thermal preference of species.

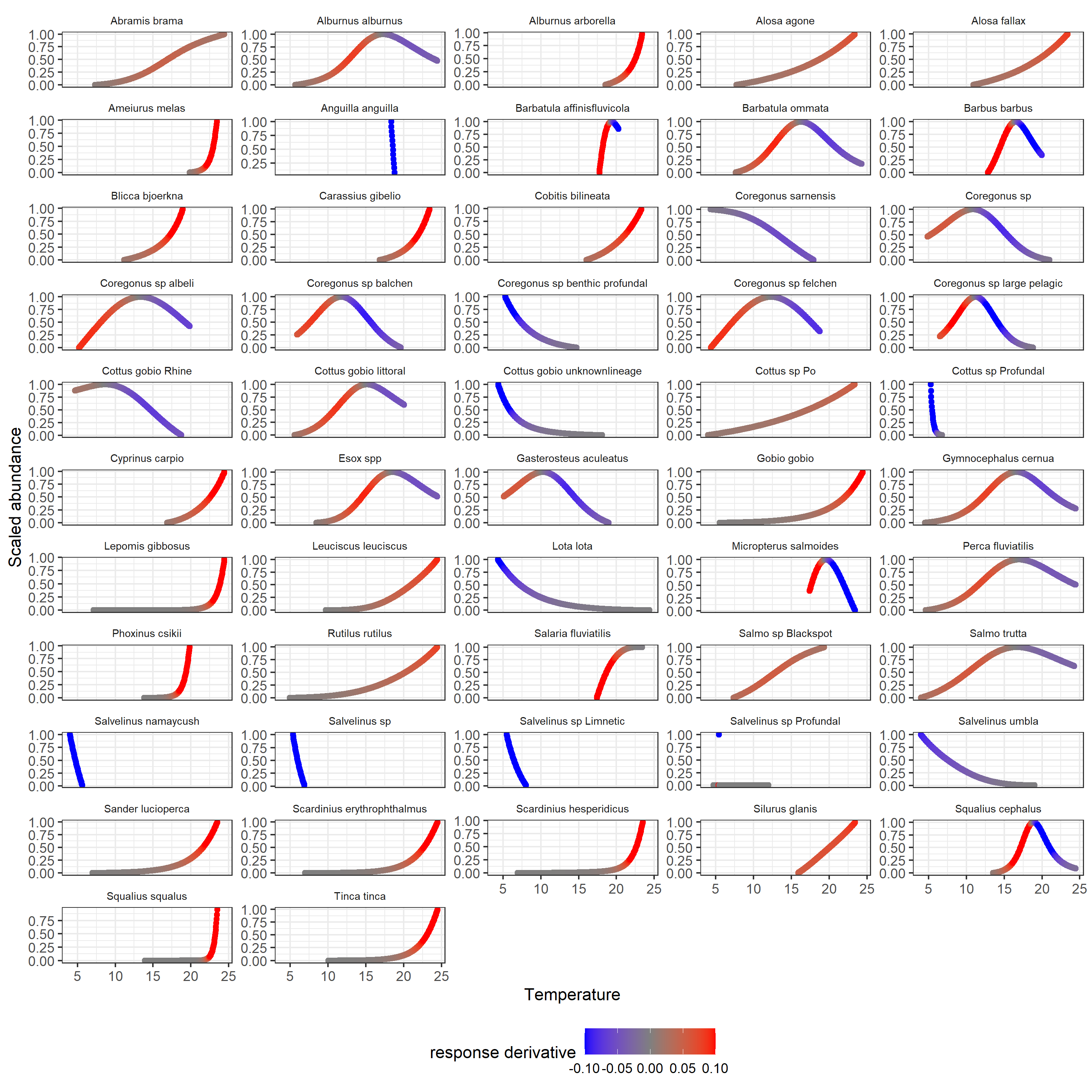

Figure S8. Scaled abundance-temperature relationships for 52 species coloured by the value of the response derivative (forward finite difference) to indicate the values contributing the response diversity in the community level metrics.

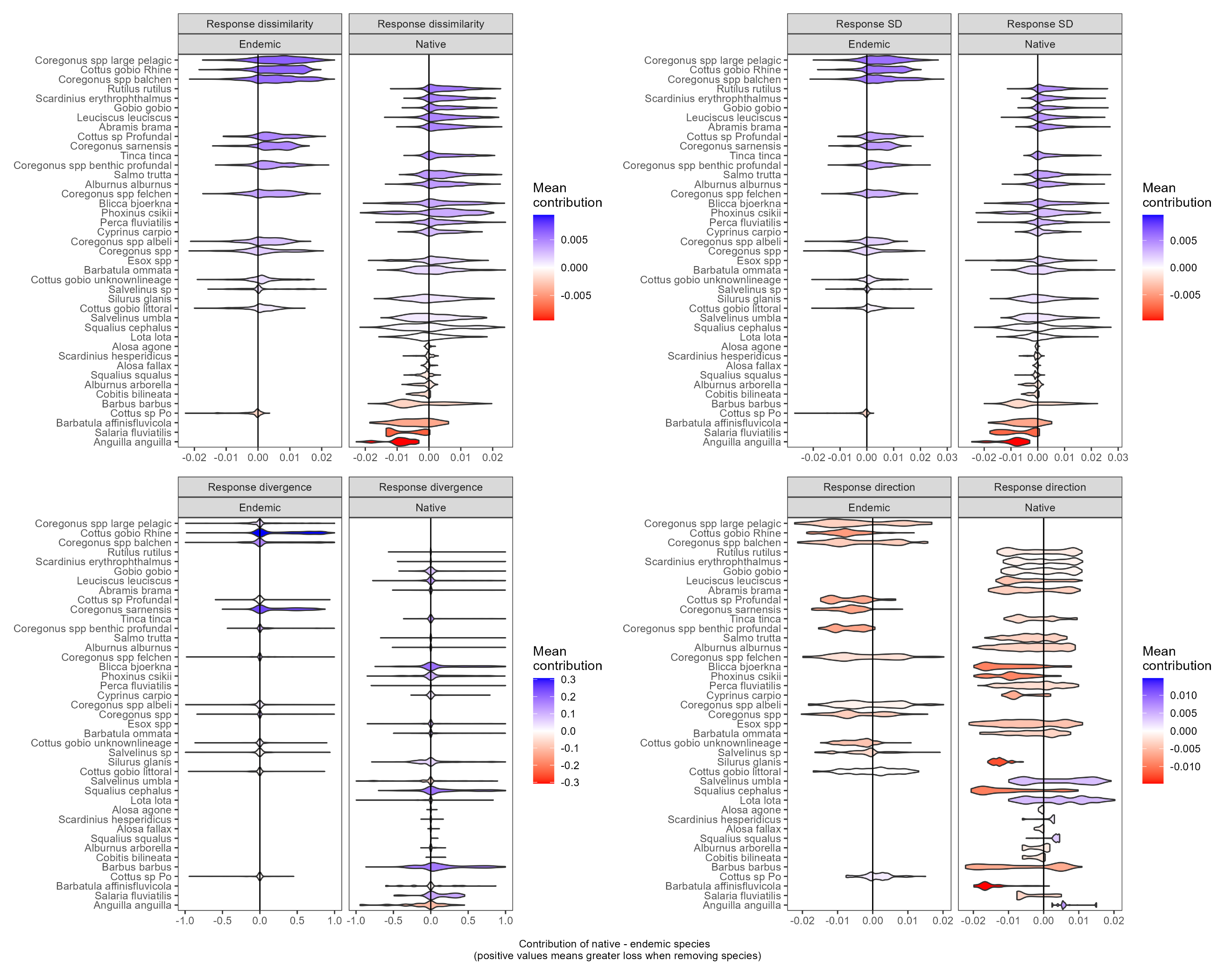

Figure S9. Difference in response diversity between endemic and native species removal from a standardized community of 10 species across 11 lakes and 10 replicates. Violins are not aggregated across simulations but show raw distributions of all simulated comparisons. Full statistical results are reported in Table S3.

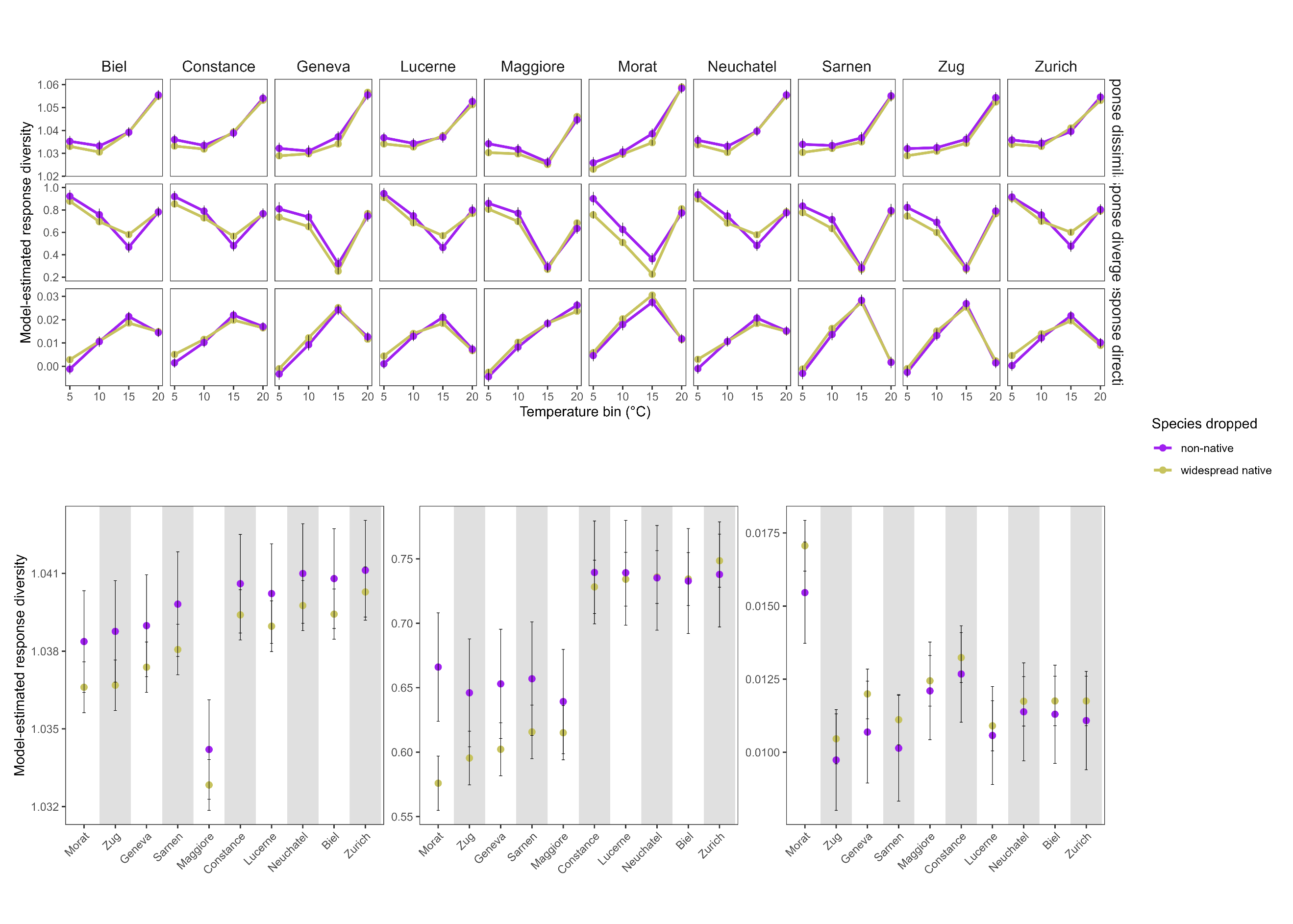

Figure S10. Species biogeographic contributions to response diversity across thermal gradients and lakes as indicated by a dropping one random species in each focal group from a standardized community. Estimated marginal mean values ± 95% CI from an ANOVA design shows the effect of each grouping, thermal strata and lake on response diversity metrics as indicated on the y-axis. The panels a-c show estimate values across temperature strata. The panels d-f show estimates across each lake. Colours indicate when in-situ diversification (green) are dropped from the community compared with widespread native species (yellow).

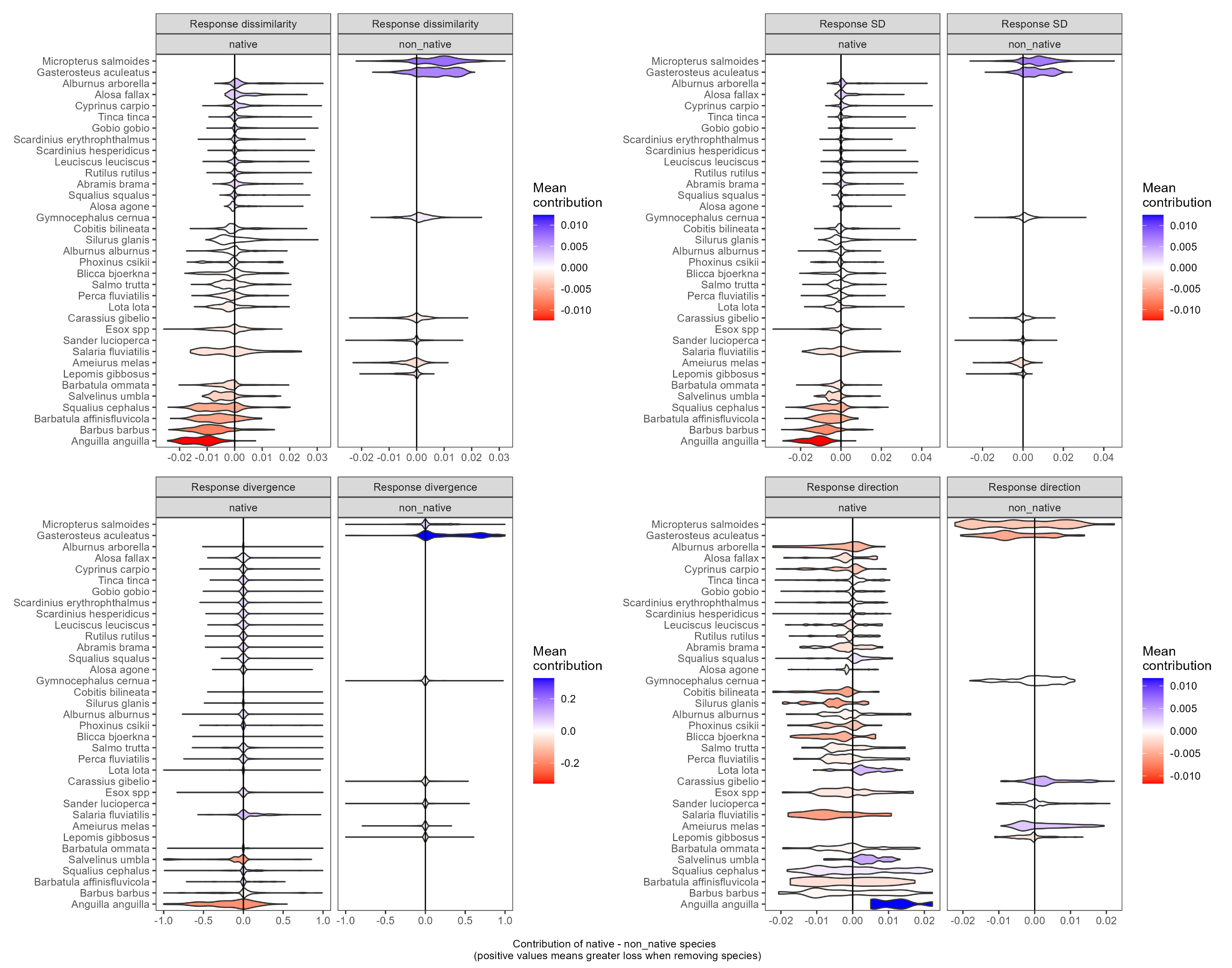

Figure S11. Difference in response diversity between non-native and native species removal from a standardized community of 10 species across 11 lakes and 10 replicates. Violins are not aggregated across simulations but show raw distributions of all simulated comparisons. Full statistical results are reported in Table S3.
