## Supporting Information 2 for "Anthropogenic-driven loss of an adaptive radiation reduces thermal response diversity"

Supplementary methods

**Details of fish community surveys**

We analysed standardized sampling of fish communities from data collected during “*Projet Lac*”, a large project that assessed fish diversity in peri-alpine lakes (Alexander & Seehausen, 2021; Figure S1). Of the 35 lakes sampled, we analysed 16 lakes where corresponding 1D temperature data were available alongside biodiversity data at project initiation in 2019 (see below; (Gaudard *et al.* 2019)). Fish communities were sampled in late summer to early autumn using gillnets (European Standard benthic and pelagic gill nets (CEN-EN 14757) and vertical gill net protocols) and electro-fishing (Figure S2). The sampling protocols provided standardized sampling across the full depth range of large lakes in Switzerland and were designed to standardise effort based on lake size (for full details see Alexander & Seehausen, 2021). The individuals caught in one set of nets or electrofishing section defined a single observation for a location. We standardized net fishing events per duration of soaking, giving catch per unit effort per 14-hour period. Electrofishing abundance estimates remained unstandardized and so we accounted for differences in methodological aspects of our sampling in our statistical approach (see below). We estimated the depth of the benthic sampling net/electrofishing event taking the mean of the maximum depth and minimum depth minus 1.5 meters to account for net height. To ensure more consistent matching between sampling and temperature data, we removed all sampling events of benthic nets with a depth range greater than 5 meters. We used the distance from the surface for pelagic nets. Note that the CEN pelagic nets were 6m heigh and we used the middle of this net as the approximate of our sampled depth estimate. In total, the data consisted of 4,239 unique observations of 52 fish species among the 16 lakes (Figure S1). On average, there were 265 sampling events per lake ranging from 98 in the smallest lake (Joux) to 638 in the largest lake (Geneva; Table S1). In total, there were 11,449 observations of species abundance from which to build models, and each species was present in an average of 220 sampling events ranging from 3 (*Salvelinus namaycush*) to 3119 (*Perca fluviatilis*; summarised in Table S2). We removed 19 species that were associated with lotic environments that had fewer than 10 sampling presences as they were both rarely encountered and not the focus of this work.

The fish taxonomy of Switzerland and its lakes has been updated in the past decade. As such, fish were first identified directly in the field following Kottelat & Freyhof (2007) and most were independently re-examined after the fieldwork based on photographs and preserved specimens, especially taxonomically challenging groups such as *Coregonus* (Selz *et al.* 2020, 2025; Selz & Seehausen 2023). Across Swiss lakes the genera *Coregonus* and *Salvelinus* have undergone adaptive radiations with distinct species between lakes having convergent ecomorphological, habitat and functional characteristics (Alexander & Seehausen 2021; De-Kayne *et al.* 2022; Vonlanthen *et al.* 2012). Infrequently caught species with similar ecomorphology were pooled into species groups in our analysis (see Table S4). For the *Coregonus* complex, we separated five ecomorphs based on spawning depth, prey type and gill raker count as: “Large pelagic”, “Benthic profundal”, “Felchen”, “Albeli” and “Balchen” (De-Kayne *et al.* 2022; Hudson *et al.* 2011). We retained *Coregonus* sp*.* for individuals not identified to species level. In Lake Sarnen, we kept *Coregonus sarnensis* as it does not fit with the other ecomorph classifications. For the five lakes with radiations in *Salvelinus* we used two ecomorphs (profundal and limnetic) and retained *S. umbla* and *S. namaycush* where present*.* We pooled all deep-water Cottus as a profundal ecomorph across lakes, and shallow water populations as a littoral ecomorph as largely supported by distinct depth distributions, genetic structure and geometric morphometrics. Finally, the genus Esox has different species north and south of the alps, but the southern species (*Esox cisalpinus*) had very few records, so we pooled these records with *Esox lucius* to enable more robust modelling (i.e., assuming niche conservatism).

We defined three biogeographic groups of species as “*widespread native”* (n=29) which occur across Europe and are not endemic to Switzerland, “*in-situ diversification” (n=15)* as those from the genera Coregonus, Salvelinus and Cottus which arose through post-glacial adaptive divergence to novel ecological opportunities in deep lake ecosystems starting around 12,000 years ago, and “*non-native*” (n=8) species are introduced to Switzerland. Table S4 gives an overview of nomenclature and biogeographic classification of species.

**Modelling species’ temperature response curves**

We quantified species’ realized thermal niches by modelling the response of abundance to temperature. We fitted generalized additive mixed effects models (GAMMs) for each of the 52 fish species using abundance as a response variable and temperature as a predictor (Figure S3). We fitted GAMMs with error distributions that matched species’ abundance. Specifically, we used random intercepts to account for the difference in mean abundance between sampling (i.e., survey method) and cover the differences between lakes that are not incorporated as separate covariates due to low replication of lakes (i.e., ecological differences between lakes). Species that had a maximum abundance of 1 in any sample across all lakes were modelled with a binomial distribution. Species that had a maximum abundance > 1 (i.e., count data) were modelled using a zero-inflated Poisson distribution (Zuur *et al.* 2014). If a species occurred in multiple lakes, data from all lakes were included in the species’ model and the lake identity was included as a random intercept. In all models, we also included the sampling protocol as a random intercept to account for differences in mean abundance recovered with different methods. We did not expect different methods to recover different realized thermal niche shapes, so opted for a more parsimonious model without a random slope term. We visualised the influence of adjusting the basis spline k from 2 to 10 on response shapes, with extreme values of k giving ecologically unrealistic response curves. We used a k of 3 which balanced curves non-linearity with physiologically realistic unimodal or sigmodal thermal responses (Clarke 2017). All GAMMs were modelled using the *mgcv* package (Wood 2011). We validated the fitted models using the *DHARMa* package by assessing the q-q plots and the simulated residuals plotted against the predicted values (Hartig 2026). We found 63% of the simulated residuals of species models showed no significant deviation from normality, with 98% of species models having visually non-skewed residuals. The aim of our work was primarily to characterise and compare the shape of realized thermal responses amongst species, rather than test statistical inferences of temperature as a driver of abundance, as such we progressed using all species in our response diversity calculations. On average, our models for a k=3 explained 57±19% of deviance in abundance. We performed sensitivity checks of our models by incorporating depth as an additional covariate, which helps incorporate gradients of light, pressure and oxygen in an indirect way. We compared our temperature-only models to models that also included depth as an explanatory variable of species abundance, using a k=3 for sample depth (Figure S4). While these models performed marginally better than temperature only models, we found they were often biologically implausible, so we preferred to work with the biologically plausible response curves from the temperature-only regressions to assess thermal response diversity (see supporting information 2 for further explanation).

Sensitivity testing of species’ temperature response curves

The quantification of species realized thermal niches may be influenced by gradients of additional environmental variables such as light, pressure and oxygen which generally correspond to depth. We therefore tested how incorporating depth as a proxy for these additional environmental gradients influence the realized thermal niche and response derivatives recovered. We compared our temperature-only models to models that also included depth as an explanatory variable of species abundance, using a k=3 for sample depth (Figure S4). We produced marginal predictions of abundance at each temperature by predicting at the mean depth value of the species. We compared the AICs and deviance explained of each model (Figure S5-S7). In general, models including depth and temperature had marginally higher deviance explained (~1%) and lower AIC values indicating better ability of temperature-depth models to explain abundance variation, as expected when incorporating a proxy variable for multiple co-varying environmental gradients. However, when compared with established thermal preferences of species and families, the thermal response curves from models incorporating depth were often biologically implausible. This is due to a well-established challenge with multiple regression, whereby coefficients (and response curves) can be less precisely estimated in the face of collinearity (Morrissey 2018). For instance, thermal response curves including depth indicated species having biogeographic affinities for temperate regions have realized thermal optima <10°C (*Barbatula, Barbus, Carassius, Cottus, Micropterus, Squalius, Scardinius*), or response curves that have a biological implausible “U” shape (*Lepomis*, *Gobio*). In contrast, temperature-only models generally produced realized thermal niches that align with established thermal preferences of fishes in our study region (Bonaglia *et al.* 2025; Freyhof & Kottelat 2007). Given that without experimentation we cannot remove this collinearity, and we expect temperature to be a major factor structuring lake fish assemblages and their diversity (Keller & Seehausen 2012; Magnuson *et al.* 1979), we preferred to work with the biologically plausible response curves from the temperature-only regressions to assess thermal response diversity.

Community metrics of thermal response diversity

We calculated thermal response diversity from the variation in species responses to temperature. We measured the instantaneous responses of species characterised by the first derivative of the (non-linear) slope between abundance and temperature, at a given temperature (Figure 1; following Ross et al. 2023). To assess the variation in thermal responses at a given temperature, we simulated the full temperature gradient in Swiss lakes and made predictions of abundance for each species given the temperature (Figure 1a and 1b). We calculated the change in scaled abundance (0-1) per 0.1°C change in temperature by calculating the forward finite difference (ffd) as:

$ffd= \frac{{abundance}_{j}-{abundance}_{i}}{0.1}$, where *i* indicates abundance at current temperature and *j* indicates abundance after a 0.1°C increase in temperature (Figure 1c and 1d; Figure S8). This approximation of a derivative from predicted values was a simple, fast and intuitive method that could be applied across models with different error distributions. This step size is appropriate as it is sufficiently small to achieve our estimate of non-linear abundance change per unit temperature change (i.e., ~0.3% of the thermal gradient across Swiss lakes at each step). To align with other literature, we hereafter refer to the *ffd* as “response derivatives” (e.g., (Ross *et al.* 2023)). To be conservative and avoid extreme response derivatives which bias estimates of response diversity (see below), we bounded the most extreme 10% of response derivatives to be 0.1 (the upper 90th quantile of absolute response derivative values).

From these response derivatives, we calculated three metrics of response diversity and one summary of net response direction (i.e., positive or negative net community response). From the response derivatives of each species, we calculated community scale response “dissimilarity” and “divergence” following Ross et al. (2023), as well as simply the standard deviation of response derivatives. These metrics were then calculated at a given 0.2°C temperature interval by summarising all species response derivatives at that temperature.

Dissimilarity and standard deviations quantify the overall difference in species responses. Dissimilarity indicates how differently species respond to a given environmental condition and is assessed based on a similarity-scaled metric of diversity based on the pairwise Euclidean distances in the response derivatives between all pairs of species in a community (Leinster & Cobbold 2012). We set the parameter *q,* which adjusts for relative abundance differences, to 0 such that the relative abundance was excluded in the calculation of dissimilarity The dissimilarity metric as presented by Leinster and Cobbold (2012), and summarised in Ross et al., (2023), has three relevant features: i) it cannot decrease with species additions, ii) adding redundant species responses (i.e., perfectly similar responses) does not increase the metric, and iii) it can detect even subtle differences in responses among the community. Dissimilarity is lowest (1), when all species respond identically and highest when all species respond differently. It can take its highest value as the species richness (total number of species in the community) of the community. Despite the above properties, we found the scaling of dissimilarity when applied to abundance models that predict most abundances to be < 1 unintuitive (varying between 1 and 1.1), so also included the standard deviation of response derivatives – but note that these metrics were highly correlated (Pearson’s correlation = 0.97).

Divergence captures the ability of a community to compensate negative responses with neutral or positive ones, as classically conceptualised as the “portfolio effect” (Schindler *et al.* 2015). That is, if some species respond positively while others respond negatively, the overall community abundance can be maintained by increases counteracting decreases. Divergence was calculated following Ross et al. (2023) as:

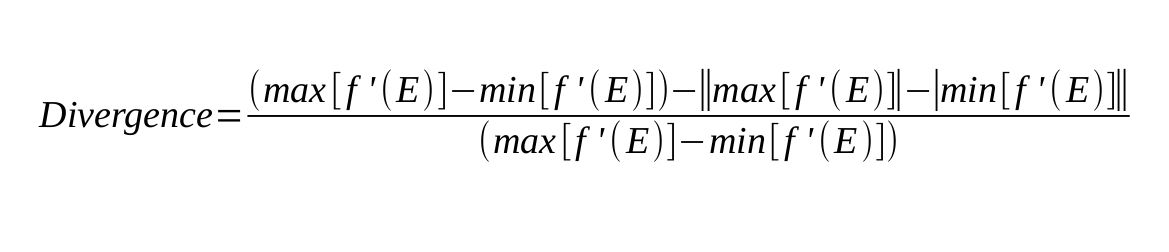

f’(E) represents the response derivative. Divergence is 0 when all response derivatives are in the same direction (i.e., all positive or all negative). If response derivatives span across zero, divergence is > 0 and < 1. Divergence takes its maximum value of 1 when the absolute values of minimum and maximum response derivatives are equal.

For each lake, we constructed potential local communities at each temperature. We used a synthesis of fish presences in Swiss lakes from Table 16 in Alexander & Seehausen (2021) to compile a complete species list for each lake. For every 0.2°C temperature bin in each lake, we then calculated the community response diversity from the set of species’ response derivatives.

**Identifying biogeographic, environmental and anthropogenic correlates of thermal response diversity**

We first assessed the patterns of metrics of response diversity and direction across thermal gradients for each lake. Next, we averaged the response diversity and direction metrics across the entire thermal gradient, giving a single response diversity and direction metric for each lake. We used spearman’s rank correlation to test how the metrics of response diversity and direction were correlated to the richness of the entire lake as well as the species richness of each biogeographic class (widespread native, in-situ diversification, non-native). Next, we assessed the same relationships but within four thermal bands within each lake (2.5 to 7.5, 7.5 to 12.5, 12.5 to 17.5 and 17.5 to 22.5) because different biogeographic groups dominate in each of these thermal bands. Finally, we assessed how historic eutrophication of lakes related to response diversity and direction in each temperature band. We used the maximum historic phosphorus levels of each lake from Vonlanthen et al. (2012). Each lake reached a different maximum total phosphorus value, with some lakes being weakly impacted and others becoming highly eutrophic. For Constance and Zurich, which each have two basins, we took the phosphorus level of the larger basin. We excluded Lake Poschiavo from this analysis as phosphorus values have been measured inconsistently and were last recorded in 1995 (Vonlanthen et al., 2014). Note that such relationships where richness and eutrophication may jointly influence overall response diversity would be best tested using multiple regressions which provide a marginal effect of each variable independently. However, we have a sample size of 16 lakes and therefore few degrees of freedom, so we avoided the risk of type II errors (false negative) by using spearman’s rank correlations and acknowledge that our inferences are open to confounding biases of richness or eutrophication in each univariate test.
